# Human-specific *NOTCH*-like genes in a region linked to neurodevelopmental disorders affect cortical neurogenesis

**DOI:** 10.1101/221226

**Authors:** Ian T Fiddes, Gerrald A Lodewijk, Meghan Mooring, Colleen M Bosworth, Adam D Ewing, Gary L Mantalas, Adam M Novak, Anouk van den Bout, Alex Bishara, Jimi L Rosenkrantz, Ryan Lorig-Roach, Andrew R Field, Maximillian Haeussler, Lotte Russo, Aparna Bhaduri, Tomasz J. Nowakowski, Alex A. Pollen, Max L. Dougherty, Xander Nuttle, Marie-Claude Addor, Simon Zwolinski, Sol Katzman, Arnold Kreigstein, Evan E. Eichler, Sofie R Salama, Frank MJ Jacobs, David Haussler

**Affiliations:** UC Santa Cruz Genomics Institute, Santa Cruz, California, United States of America; University of Amsterdam, Swammerdam Institute for Life Sciences, Amsterdam, The Netherlands; Molecular, Cell and Developmental Biology, of California Santa Cruz, Santa Cruz, California, United States of America; Department of Computer Science and Department of Medicine, Division of Hematology, Stanford University, California, USA; Howard Hughes Medical Institute, University of California Santa Cruz, Santa Cruz, California, United States of America; The Eli and Edythe Broad Center for Regeneration Medicine and Stem Cell Research at UCSF, San Francisco, USA; Department of Genome Sciences, University of Washington School of Medicine, Seattle, WA, USA; Center for Genomic Medicine, Massachusetts General Hospital, Harvard Medical School, and Broad Institute of MIT and Harvard, Boston, Massachusetts, United States of America; Service de génétique médicale, Lausanne, Switzerland; Department of Cytogenetics, Northern Genetics Service, Institute of Genetic Medicine, Newcastle upon Tyne, United Kingdom; Howard Hughes Medical Institute, University of Washington, Seattle, WA, United States of America

**Author notes:** These authors contributed equally to this work (Ian T Fiddes, Gerrald A Lodewijk). Senior authors (Sofie R Salama, Frank MJ Jacobs, David Haussler). Present address: Translational Research Institute, Princess Alexandra Hospital, Brisbane, Australia (Adam D Ewing).

## Abstract

Genetic changes causing dramatic brain size expansion in human evolution have remained elusive. Notch signaling is essential for radial glia stem cell proliferation and a determinant of neuronal number in the mammalian cortex. We find three paralogs of human-specific NOTCH2NL are highly expressed in radial glia cells. Functional analysis reveals different alleles of NOTCH2NL have varying potencies to enhance Notch signaling by interacting directly with NOTCH receptors. Consistent with a role in Notch signaling, NOTCH2NL ectopic expression delays differentiation of neuronal progenitors, while deletion accelerates differentiation. NOTCH2NL genes provide the breakpoints in typical cases of 1q21.1 distal deletion/duplication syndrome, where duplications are associated with macrocephaly and autism, and deletions with microcephaly and schizophrenia. Thus, the emergence of hominin-specific NOTCH2NL genes may have contributed to the rapid evolution of the larger hominin neocortex accompanied by loss of genomic stability at the 1q21. 1 locus and a resulting recurrent neurodevelopmental disorder.

## Introduction

The human brain is characterized by a large neocortex that forms the substrate for the development of human-specific higher cognitive functions (Lui et al., 2011; Molnar et al., 2006; Rakic, 2009), but evolutionary changes to our genome that underlie the increase in size and complexity of the human neocortex are poorly understood (Varki et al., 2008). Structural genomic variants account for 80% of human-specific base pairs and are therefore an important class of genomic regions to consider (Cheng et al., 2005). Of particular interest are loci where segmental duplications have created entirely new human-specific gene paralogs that are associated with cortical development, such as *SRGAP2C* (Dennis et al., 2012), (Charrier et al., 2012), *ARHGAP11B* (Florio et al., 2015) and *TBC1D3* (Ju et al., 2016). Interestingly, human-specific duplicated genes are often located within segmental duplications that mediate recurrent rearrangements associated with neurodevelopmental disorders (Stankiewicz and Lupski, 2010) (Florio et al., 2015), (Nuttle et al., 2016), (Popesco et al., 2006), (Dumas et al., 2012), (Dougherty et al., 2017). One region susceptible to these rearrangements lies on human chromosome band 1q21, which was involved in a large pericentric inversion involving considerable gene loss and duplication during human evolution (Szamalek et al., 2006), contains a disproportionate number of human-specific genes (O'Bleness et al., 2012), and also contains the 1q21.1 distal deletion/duplication syndrome interval (Mefford et al., 2008), (Brunetti-Pierri et al., 2008). *De novo* deletion of one copy of this 1q21.1 locus frequently leads to an abnormal reduction in brain size (microcephaly) and reciprocal duplication often results in an abnormal increase in brain size (macrocephaly), among other symptoms.

The 1q21.1 locus was not correctly assembled in the human reference genome until the most recent version, GRCh38, hampering early research of 1q21.1 syndromes. This is due to several tracts of almost identical paralogous DNA. The most recent work (Dougherty et al., 2017) hypothesized that the typical breakpoints for the distal 1q21.1 deletion/duplication syndrome lie within two of these tracts, at ~chr1:146,200,000 and chr1:148,600,000 in GRCh38, respectively, in regions of ~250 kb at 99.7% identity. We confirm this, and show that these regions contain little-studied, paralogous genes we call *NOTCH2NLA* and *NOTCH2NLB* that affect Notch signaling during human neurodevelopment. There are also atypical breakpoints for the distal 1q21.1 syndrome (Dougherty et al., 2017) that reach further almost to a third *NOTCH2NL* paralog we call *NOTCH2NLC.* We show both the typical and atypical breakpoints lead to change in NOTCH2NL copy number. This suggests that *NOTCH2NL* paralogs could contribute to the brain size and other cognitive symptoms of 1q21.1 distal deletion/duplication syndrome since Notch signaling is central to brain development, determining the timing and duration of neuronal progenitor proliferation and neuronal differentiation (Louvi and Artavanis-Tsakonas, 2006), (Kageyama et al., 2009), (Hansen et al., 2010).

We also find that *NOTCH2NL* genes are unique to the human, Neanderthal and Denisovan species among all species that have been sequenced (i.e., unique to hominin species), and originated just prior to or during the period where fossil evidence demonstrates the most rapid growth in the size of the hominin neocortex (Holloway et al., 2004). *NOTCH2NL’s* origin appears to be in an ectopic gene conversion event 3-4 million years ago in which part of *NOTCH2* “overwrote” an earlier NOTCH2-like pseudogene creating a new functional, short, secreted form of *NOTCH2*.

*NOTCH2NL* is expressed in the developing fetal brain at highest levels in radial glia neural stem cells, including outer radial glia cells, a cell type hypothesized to generate the majority of primate cortical neurons and to contribute more to cortical expansion in hominin than in other apes (Lewitus et al., 2013), (Smart et al., 2002). Functional analysis of *NOTCH2NL* by ectopic expression in mouse embryonic stem cell (mESC)-derived cortical organoids, as well as analysis of CRISPR-generated *NOTCH2NL* deletion mutant human embryonic stem cell (hESC)-derived cortical organoids, suggests that *NOTCH2NL* expression delays differentiation of neuronal progenitor cells including dorsal radial glial cells. This is consistent with cerebral neoteny observed in hominin brain development where, perhaps for obstetrical reasons, the human and Neanderthal brains are slower to mature during fetal development and expand by a factor of 3.3 after birth, compared to a factor of 2.5 in chimpanzees (DeSilva and Lesnik, 2006; Ponce de Leon et al., 2008).

Finally, we demonstrate that NOTCH2NL physically interacts with NOTCH receptors and can activate NOTCH signaling in reporter assays in the presence and absence of NOTCH ligands. From functional and genomic evolutionary evidence, we propose that with the creation of the modern forms of *NOTCH2NL* genes in the last few million years, hominins gained a new, secreted NOTCH-like protein that can act to amplify Notch signaling and promote increased cortical neurogenesis by delaying differentiation of neuronal progenitor cells. Thus, the emergence of *NOTCH2NL* genes in hominins may have contributed to the increase in size and complexity of the human neocortex at the expense of susceptibility to 1q21.1 distal duplication/deletion syndrome.

## Results

### NOTCH2NL is a novel NOTCH-like gene, uniquely expressed in humans

The gene annotated as *NOTCH2NL* (Duan et al., 2004) on human genome assembly GRCh37 resides on the q arm of human chromosome 1 in the 1q21.1 locus, which long remained one of the most difficult regions in our genome to assemble due to its highly repetitive nature (Szamalek et al., 2006), (Dennis et al., 2012), (Doggett et al., 2006).The 1q21.1 locus has undergone a number of human lineage-specific rearrangements, including a pericentric inversion (Szamalek et al., 2006) and the creation of new human-specific gene paralogs, including *HYDIN2* (Dougherty et al., 2017) and *SRGAP2B* (Dennis et al., 2012). Resequencing of the pericentric region of chromosome 1 in a haploid human cell line finally resolved previously unmapped regions and led to a revised assembly of the 1q21.1 locus, which is incorporated in the human genome assembly GRCh38 (Steinberg et al., 2014). This improved assembly reveals the presence of four paralogous NOTCH2NL-like genes (**Figure 1A**): *NOTCH2NLA, NOTCH2NLB* and *NOTCH2NLC* reside in the 1 q21.1 locus, and a fourth quite different paralog, *NOTCH2NLR* (NOTCH2NL-Related) is located near *NOTCH2* on the p-arm of chromosome 1. The greater than 100 kb genomic regions spanning each *NOTCH2NL* gene show greater than 99.1% identity to *NOTCH2* (Figure S1A), suggesting that *NOTCH2NL* paralogs were created within the last few million years, on a time scale similar to that of *SRGAP2* and *HYDIN2* (Dennis et al., 2012), (Dougherty et al., 2017).

**Figure 1.**
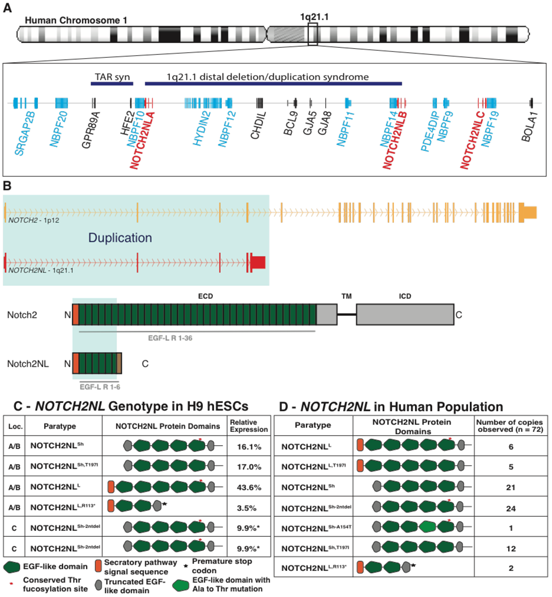
*NOTCH2NL* genes are associated with a neurodevelopmental disease locus and exhibit variable protein structures in the human population. (A) Location of *NOTCH2NL* genes in chromosome 1q21.1 flanking the 1q21.1 distal deletion/duplication syndrome locus. This region contains many additional genes derived from human segmental duplication (shaded light blue). TAR syn is Thrombocytopenia Absent Radius syndrome. (B) Gene and protein features of NOTCH2 and NOTCH2NL. (C) Results of the *de novo* assembly of *NOTCH2NL* loci for H9 human embryonic stem cells and relative allele expression from week 5 cortical organoids as measured by full-length cDNA sequencing. * Not enough nucleotide differences present to distinguish between the two *NOTCH2NL^sh2ntdel^* alleles, so counts evenly split between the two. (D) Observed *NOTCH2NL* paratypes (n=72, counting paratypes transmitted from parent to child only once) obtained from linked-read sequencing and assembly of 15 individuals. See also Fig. S1 and Table S1 and S4.

*NOTCH2NL* results from a partial duplication of the *NOTCH2* gene. The duplicated segment includes the upstream region of *NOTCH2,* the *NOTCH2* promoter and EGF domains from *NOTCH2* exons 1 through 4, while excluding the membrane spanning and cytoplasmic domains of *NOTCH2.* Because it lacks a transmembrane domain, the NOTCH2NL protein can be secreted by the cells that express it (Duan et al., 2004). *NOTCH2NL* genes contain a fifth exon derived from *NOTCH2* intronic sequence that provides NOTCH2NL with 20 unique amino acids (**Figure 1B, Figure S1I**). In paralogs *NOTCH2NLA, NOTCH2NLB* and *NOTCH2NLC* the 5th exon has a 4 bp deletion compared to the corresponding sequence in *NOTCH2*. We established that this 4 bp deletion is essential for NOTCH2NL protein expression (**Figure S1B-E**). *NOTCH2NLR* lacks this 4 bp deletion, contains many variants in its protein coding region relative to *NOTCH2* and *NOTCH2NL* paralogs including a very recent still segregating variant (**Figure S1I**), and analysis of human genome sequence data from the Simons diversity project (N=266) (Mallick et al., 2016) reveal a significant fraction of the population (14%) lack *NOTCH2NLR* (**Figure. S1G**). Together these results suggest that *NOTCH2NLR* is a non-functional pseudogene, although additional investigation would be needed to verify that its mRNA and/or protein product is not functional.

Despite the overall high sequence homology of the *NOTCH2NL-gene* loci as present on GRCh38, each *NOTCH2NL* gene paralog has distinguishing characteristics (**Figure 1D**): *GRCh38NOTCH2NLA* has an ATG→ ATA mutation in the *NOTCH2* initiator methionine codon and as a result, GRCh38NOTCH2NLA protein lacks the N-terminal 39 amino acids including the signal peptide of NOTCH2. We term this allele “short NOTCH2NL” and denote it *NOTCH2NL^Sh^. GRCh38NOTCH2NLB* encodes a longer protein with the signal peptide, but has a Thr→Ile substitution in the 5th EGF repeat, and is referred to as “long NOTCH2NL with Thr→Ile substitution” *(NOTCH2NL^L,T1971^,* coordinates relative to the NOTCH2 protein). This substitution occurs at a site for fucosylation of NOTCH2. *GRCh38NOTCH2NLC* contains a 2 bp deletion just downstream the original NOTCH2 start codon *(NOTCH2NL^Sh-2ntdel^*), and like *NOTCH2NL^Sh^*, NOTCH2NLC protein is initiated at a downstream ATG, resulting in a short NOTCH2NL protein that lacks the N-terminal signal peptide.

To better understand the spectrum of *NOTCH2NL* alleles in the human population we sought to fully resolve the *NOTCH2NL* haplotypes in several individuals. To overcome the challenge of accurately assembling these highly homologous genes we took advantage of the linked reads of readset genome sequence technology whereby small DNA molecules compatible with short read DNA sequencers are labeled (barcoded) according to individual long genomic DNA fragments (>30 kb) from which they are derived. We developed an assembly-by-phasing approach called Gordian Assembler utilizing this barcoded data to resolve the five NOTCH2-related gene paralogs in 8 normal individuals (STAR Methods, **Figure S1H, Table S1**). This and related analysis revealed recent, likely ongoing, ectopic gene conversion occurring between *NOTCH2NLA, B* and *C* (**Figure S1F**) that is so extensive that *NOTCH2NLA* and *NOTCH2NLB* act essentially as a single gene with 4 alleles, and to some extent all three paralogs act as a single gene with 6 alleles. Our discovery of a large number of gene conversion alleles segregating in the human population is consistent with previous analysis of 1q21.1 (Nuttle et al., 2013). This, coupled with ~250 kb of DNA at 99.7% identity between the genomic regions containing *NOTCH2NLA* and *NOTCH2NLB,* makes it nearly impossible to assign individual alleles to a specific locus. Focusing on one assembly, that of the H9 hESC line, we identified six different alleles derived from *NOTCH2NLA, NOTCH2NLB* and *NOTCH2NLC* (**Figure 1C, Figure S1F**). These include three additional *NOTCH2NL* alleles beyond those present in the GRCh38 reference genome, including a short version with Thr→lle (*NOTCH2NL^ShJ1971^*) and a long version without Thr→Ile (*NOTCH2NL^L^*). We also observed an additional long version with a SNP (rs140871032) causing an in-frame stop codon at amino acid 113 (*NOTCH2NL^L,R113*^*), also seen in 35 of 266 genomes in the Simons Diversity collection (Mallick et al., 2016). Finally, we observed a rare paratype *NOTCH2NL^Sh,A154T^* (rs76765512) with a non-synonymous change in EGF repeat 3 in one assembled genome and six Simons Diversity collection genomes. By RNA-seq analysis of cortical organoids derived from H9 hESCs (see below), we confirmed that all the *NOTCH2NL* alleles assembled in H9 are expressed. *NOTCH2NL^L,R113*^* represents only about 3.5% of the transcripts, perhaps due to nonsense-mediated decay (**Figure 1C**). In the 17 genomes in which we have determined 72 *NOTCH2NL* alleles, we have found seven different *NOTCH2NL* sequence variants that affect the features of the resulting protein (**Figure 1D, Figure S1I**). We refer to these various *NOTCH2NL* alleles as paratypes since their physical location can vary among paralogous locations and therefore they do not conform to standard haplotypes. A typical *NOTCH2NL* genotype consists of 6 *NOTCH2NL* paratypes (**Figure 1C**), rather than two haplotypes as for most genes. Sequencing of these initial 72 NOTCH2NL-related paratypes is unlikely to have revealed all the structural variation in *NOTCH2NL* in the human population. We expect more variants will be found as additional genomes are sequenced and assembled in a manner allowing us to read these paratypes (**Figure 1D, Table S1, S4**).

### Multiple rounds of gene duplication and gene conversion lead to functional *NOTCH2NL* genes only in hominins

To explore the evolutionary history of *NOTCH2NL* genes in more detail, we assessed the presence and structure of *NOTCH2NL* genes in other primate genomes. Based on alignment of genomic DNA-reads matching or nearly matching the parental *NOTCH2* locus, and determination of allele frequency distributions of Singly Unique Nucleotides (SUNs), base variants that occur only in one paralog (Sudmant et al., 2010), we established that the *NOTCH2NL* sequence emerged by a partial duplication of *NOTCH2* prior to the last common ancestor (LCA) of human, chimpanzee and gorilla. Based on absence in orangutan and all other outgroups, it occurred after the LCA of human, chimpanzee, gorilla and orangutan (**Figure 2**). Both chimpanzee and gorilla have variable read depth over the region encompassing their *NOTCH2NL*-like sequences, suggesting they contain multiple versions of truncated *NOTCH2* putative pseudogenes sharing the same breakpoint within *NOTCH2* present in the human *NOTCH2NL* genes (**Figure 2A**). Using Bacterial Artificial Chromosome (BAC) clones and whole-genome shotgun contigs, we identified several individual *NOTCH2NL*-like genes in the chimpanzee genome, all of which were truncated and predicted to be non-functional (**Figure 2B, Figure S2**). Some of these chimpanzee *NOTCH2NL*-like pseudogenes lack a 52 kb region containing exon 2. The resulting *NOTCH2NL*-like Aexon2 transcripts encode a protein of 88 amino acids with no homology to human NOTCH2NL (**Figure S2F**), which was confirmed by ectopic expression of one of these chimp *NOTCH2NL*-like cDNAs (**Figure S2D**). From whole-genome shotgun contigs we found evidence for additional *NOTCH2NL*-like pseudogenes in the chimpanzee lacking either exon 1 or exon 1-2, located downstream of various 3’ truncated genes (**Figure 2B, Figure S2B**) These make pseudogene fusion-transcripts of the form *PDE4DIP^exon1-27^-NOTCH2NL^exon2-5^, TXNIP^exon1^-NOTCH2NL^exon2-5^ MAGI3^exon1^-NOTCH2NL^exon3-5^,* and *MAGI3^exon1-14^-NOTCH2NL^exon3-5^* as confirmed by RT-PCR using RNA derived from chimpanzee induced pluripotent stem cells (IPSCs) (**Figure S2C**). Sequence analysis of these transcripts established that the *NOTCH2NL* exons in all of these fusion-genes were out of frame with the upstream exons, indicating that these fusion genes cannot produce functional NOTCH2NL proteins (**Figure S2F**). In support of this result, whereas we can detect NOTCH2 protein in extracts derived from chimpanzee IPSCs, we cannot detect a NOTCH2 N-terminal antibody reactivity in the molecular weight range expected for NOTCH2NL proteins under the same conditions where we can detect human NOTCH2NL protein (**Figure 2C**). Overall, the analysis of the configuration of *NOTCH2NL*-like genes in the chimpanzee genome and experimental evidence of lack of the protein indicates that chimpanzees lack a functional *NOTCH2NL* gene.

**Figure 2.**
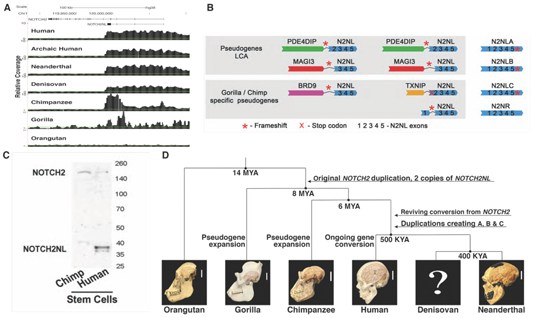
Evolutionary analysis of **NOTCH2NL*-like* genes reveals only hominin *NOTCH2NL* genes encode NOTCH-related proteins. (A) Coverage of Illumina genome sequencing reads mapped to the *NOTCH2* locus demonstrating an excess of coverage in the 5’ region of *NOTCH2* into intron 4 in all hominins examined, chimpanzee and gorilla, but not orangutan. (B) Schematic of NOTCH2NL-containing genes identified in gorilla, chimpanzee and human demonstrating that only in human *NOTCH2NL* genes encode NOTCH2-related proteins. (C) Immunoblot using an N-terminal NOTCH2 antibody (aa 25-255) of protein extracts from human and chimpanzee pluripotent stem cells showing expression of full-length NOTCH2 in both samples, but NOTCH2NL only in human. (D) Summary of the major events leading to the emergence of *NOTCH2NL-related* genes in the great ape lineage. See also Figure S2.

In the gorilla genome, three fusion pseudogenes involving *NOTCH2NL*-like sequence were identified (**Figure S2A**). In one gorilla BAC, exons 1-12 of *BRD9* are fused to exons 3-5 of NOTCH2NL. Surprisingly, two other gorilla fusion pseudogenes were similar to NOTCH2NL-fusion pseudogenes found in chimpanzee: a *PDE4DIP^exon1-27^-NOTCH2NL^exon2-5^* fusion was supported by BAC sequence and a *MAGI3^exon1-14^-NOTCH2NL^exon3-5^* fusion was supported by trace archive sequence. Transcript support was obtained for *BRD9-NOTCH2NL* and *PDE4DIP^exon1-27^-NOTCH2NL^exon2-5^* using RNA from gorilla iPSCs (Marchetto et al., 2013) (**Figure S2C**).

The presence of two distinct fusion pseudogenes in both gorilla and chimpanzee, *PDE4DIP-NOTCH2NL* and *MAGI3-NOTCH2NL*, suggests that these two pseudogenes were established in the LCA of human, chimp and gorilla. Yet none of the chimpanzee or gorilla pseudogenes are found in the human genome. Further, human *NOTCH2NL* genes are all in the vicinity of *PDE4DIP,* but in stark contrast to the *PDE4DIP-NOTCH2NL* fusion genes in chimp and gorilla, human *NOTCH2NL* genes have a 5’ genomic structure highly similar to *NOTCH2.* This suggests a plausible evolutionary history of *NOTCH2NL* genes as follows (**Figure 2D**): Both the *PDE4DIP-NOTCH2NL* and *MAGI3-NOTCH2NL* fusion pseudogenes were present the common ancestor of human, chimp, and gorilla between 8 and 14 MYA. Then, only in the human lineage, the ancestral *PDE4DIP-NOTCH2NL* fusion gene was ‘revived’ by *NOTCH2* through ectopic gene conversion. With the acquisition of exon 1 and the upstream promoter, this created a viable *NOTCH2NL* gene encoding a stable NOTCH-related protein. Because no remnants of the *MAGI3-NOTCH2NL* fusion pseudogene are found in the human genome, this must have been lost, perhaps in the upheaval of the pericentric inversion on chromosome 1 and subsequent large-scale copy number changes (Szamalek et al., 2006). The revived human *NOTCH2NL* subsequently duplicated twice more to form the cluster of three nearly identical *NOTCH2NL* genes on the q arm of chromosome 1 in the human genome. Chimpanzee and gorilla had additional, species-specific *NOTCH2NL-*like gene duplications, but none produced functional genes.

It should be noted that ectopic gene conversion is not unusual in this region (Nuttle et al., 2013). This scenario involves quite a number of lineage-specific gene duplications, rearrangements and other losses. Close neighbors of *NOTCH2NL* have a similarly elevated number of lineage-specific structural events, e.g., the *NBPF* paralogs, which are estimated to have had six lineage-specific duplications in the human lineage (O'Bleness et al., 2014). This propensity for segmental duplication, loss, rearrangement and ectopic gene conversion is characteristic of what have been called duplication hubs in the ape genomes (Bailey and Eichler, 2006). *NOTCH2NL* is part of such a hub.

To explore when the gene conversion event that revived *NOTCH2NL* occurred, we calculated the number of substitutions per kilobase between *NOTCH2* and *NOTCH2NL,* and we analyzed three archaic human genomes (Lazaridis et al., 2014), two Neanderthal genomes (Prufer et al., 2017), (Prufer et al., 2014) and one Denisovan genome (Meyer et al., 2012). While archaic humans appear to have three *NOTCH2NL* loci (presumably A, B and C) and one *NOTCH2NLR* like modern humans, the Neanderthal and Denisovan genomes have only A, B and C-like sequence, and lack *NOTCH2NLR.* The presence of *NOTCH2NLA, B,* and *C* in all 3 hominin genomes suggests that the ectopic gene conversion creating hominin *NOTCH2NL* happened prior to their LCA, more than 0.5 MYA. The substitution rate between human *NOTCH2* and human paralogs *NOTCH2NLA, B* or *C* is roughly half of that between human and chimp, which, if calibrated to a human-chimp divergence of 6.5 MYA, gives a date for the ectopic gene conversion between 3 and 4 MYA. This corresponds to a time just before or during the early stages of the expansion of the hominin neocortex (Holloway et al., 2004) (**Figure S2E**).

### NOTCH2NL is expressed in Radial Glia neural stem cells during human cortical development

*NOTCH2NL* shares promoter sequence with *NOTCH2* and is therefore expected to be expressed in a pattern similar to *NOTCH2.* To learn what cell types in the developing brain express *NOTCH2NL,* we examined *NOTCH2NL* expression in 3,466 single cells derived from human fetal brains ranging in age from 11 to 21.5 post coital weeks (pcw) that were sampled from multiple regions of the dorsal and ventral telencephalon (Nowakowski, et al, *Science,* In Press, <bit.ly/cortexSingleCell>) (**Figure 3**).

**Figure 3.**
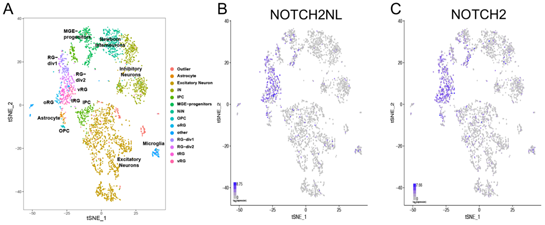
Radial glia-specific expression of *NOTCH2NL* in human fetal brain samples. Scatterplot of 3466 fetal brain cells after principal components analysis and t-stochastic neighbor embedding (tSNE) samples as described in Nowakowski, et al, 2017. Cells are colored by annotated cell type clusters (A), *NOTCH2NL* expression (B) and *NOTCH2* expression (C). See also Figure S3.

This data set contains a broad diversity of cell types including radial glia neural stem cells, intermediate progenitor cells (IPC), excitatory and inhibitory neurons, oligodendrocyte progenitors (OPC), astrocytes and microglia. The *NOTCH2NL* expression pattern we found closely resembles that of *NOTCH2* and is highest in various radial glia populations, including outer radial glia cells (oRG), as well as astrocytes and microglia (**Figure 3, Figure S3**). *NOTCH2NL* expression in the oRG population is especially interesting as this pool of neural stem cells is thought to be responsible for much of the extended neurogenesis unique to human. This neurogenesis occurs in the more sparsely packed outer subventricular zone and recent transcriptional analysis suggests that oRG cells themselves express factors to support the subventricular niche (Pollen et al., 2015). In this setting, a secreted factor like NOTCH2NL might have a critical role in promoting maintenance of the neural stem cell fate.

We also estimated the relative expression of the H9 hESC paratypes of *NOTCH2NL* in bulk RNA undifferentiated cells and week 5 cortical organoids derived from H9 ESCs (see below). We used two different methods for this analysis. First, we measured the relative abundance using paratype-specific features in Illumina short-read RNA-seq data (**Figure S3B**). This analysis suggests that *NOTCH2NL^L^* has the highest expression, and the other *NOTCH2NL* paratypes are expressed at levels 20-60% of *NOTCH2NL^L^.* These estimates have some degree of uncertainty as much of the short read data is uninformative as to which paratype is being measured. As an alternative approach, we made a full-length cDNA library enriched for transcripts originating from chromosome 1q21.1 using biotinylated oligos and sequenced this library using a MinlON nanopore sequencer. The resulting full-length *NOTCH2NL* transcripts were then assigned to their appropriate paratype, which allowed for a more accurate estimation of relative paratype expression (STAR Methods). The results confirmed that *NOTCH2NL^L^* has the highest expression, accounting for 43.6% of the transcripts, and indicated that the other paratypes have expression levels between ~8% and 40% of *NOTCH2NL^L^* (**Figure 1C**).

### Ectopic expression of NOTCH2NL delays mouse cortical neuron differentiation

To address the role of *NOTCH2NL* in cortical development, we assessed the effects of ectopic NOTCH2NL in mouse cortical organoids. We generated stable cell lines of mouse ESCs ectopically expressing human *NOTCH2NL^Sh,T1971^*, and mESC EV control cell lines. mESCs were differentiated into cortical organoids using an established protocol (Eiraku et al., 2008) (**Figure 4A**).

**Figure 4.**
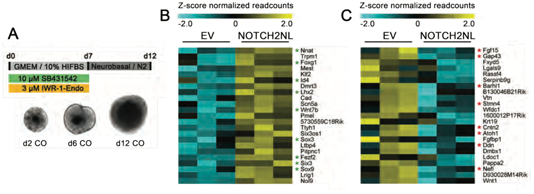
Ectopic expression of *NOTCH2NL* delays neuronal differentiation. (A) Overview of mouse cortical organoid differentiation protocol. (B) Genes upregulated in mouse organoids ectopically expressing *NOTCH2NL^Sh,T197I^* compared to EV. Asterisks indicate genes associated with radial glia cells. (C) Genes downregulated in mouse organoids ectopically expressing *NOTCH2NL^Sh,T197I^* compared to EV. Asterisks indicate genes involved in neuron differentiation. n = 3 per condition, pools of 16 organoids per replicate. See also Figure S4 and Table S2.

At day 6 of differentiation, a stage where most cells express the neural stem cell marker Pax6, mouse cortical organoids were harvested for RNA sequencing (RNA-seq). Differential expression analysis identified 147 differentially expressed genes (p-adj < 0. 05, **Figure S4A-B, Table S2**). Mouse organoids ectopically expressing NOTCH2NL showed increased expression of genes involved in negative regulation of neuron differentiation, such as *Foxgl, Id4, Fezf2, Sox3* and *Six3* (**Figure 4B**) and several genes associated with neuronal differentiation were downregulated, including *Cntn2, Nefl, Gap43,* and *Sox10* (**Figure 4C, Figure S4C**). These results suggest ectopic expression of NOTCH2NL in mouse organoids delays differentiation of neuronal progenitor cells.

### Deletion of NOTCH2NL affects development of human cortical organoids

To explore the functional role of *NOTCH2NL* in human cortical development, we used the CRISPR/Cas9 system to generate genetic deletions of *NOTCH2NL* genes in hESCs. To avoid targeting *NOTCH2*, whose sequence is >99% identical to *NOTCH2NL* even in intronic regions, we used two guides, one in intron 1 with a 1 base mismatch with *NOTCH2* and *NOTCH2NLR,* but identical to the corresponding sequence in all H9 1q21 *NOTCH2NL* genes, and another that spans a 4 bp deletion relative to *NOTCH2* at the start of exon 5. This region is also quite different in *NOTCH2NLR* (13/20 mismatches to *NOTCH2NL*) (**Figure S5A**).

Clones were screened by PCR for the large deletion from intron 1 to exon 5 and then several were expanded and analyzed by targeted linked-read sequencing and assembly as described above to determine which NOTCH2NL genes were affected and to evaluate potential off-target effects at *NOTCH2.* For one clone, the sequence analysis revealed that *NOTCH2NLA* and *B* genes showed a homozygous deletion, and NOTCH2NLC a heterozygous deletion, leaving only one NOTCH2NLC (*NOTCH2NL^Sh-2ntdel^)* intact. This clone is denoted H9*^NOTCH2NLΔ^* (**Figure 5A-B**). As a control, another clone was selected that went through the same CRISPR/Cas9 transfection and selection process, but does not harbor any alterations around the guide sites in *NOTCH2NL, NOTCH2NLR* or *NOTCH2* loci (denoted H9*).

**Figure 5.**
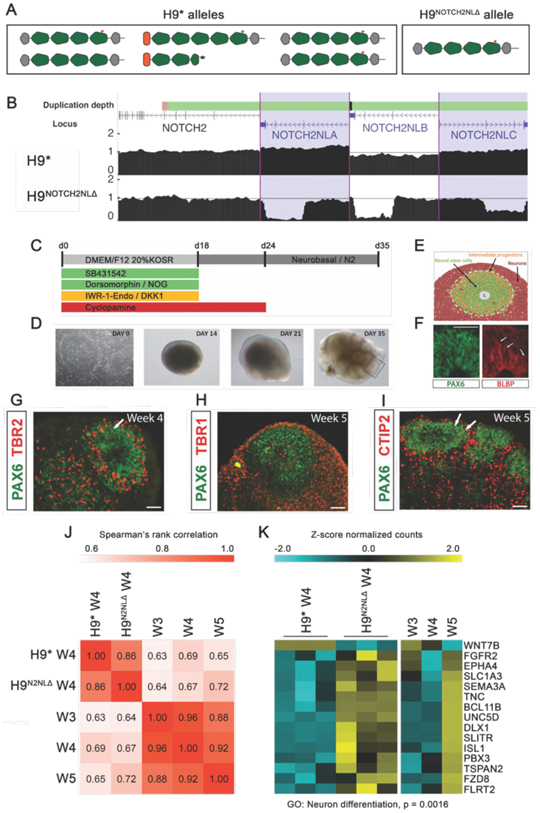
Cortical organoids from hESCs lacking *NOTCH2NL* show premature neuronal maturation. (A)Schematic showing *NOTCH2NL* alleles present in control (H9*) and NOTCH2NL mutant (H9^NOTCH2NLΔ^) cell lines. (B). Multi-region UCSC genome browser view of *NOTCH2* and *NOTCH2NL* genes with tracks showing normalized genome sequencing coverage demonstrating homozygous loss of exon 2-5 sequence for *NOTCH2NLA* and
*NOTCH2NLB* and heterozygous loss for *NOTCH2NLC*. Some coverage is seen in *NOTCH2NLA* and *NOTCH2NLB* due to a small portion of ambiguous linked read barcodes. Schematics of the cortical organoid protocol used (C) with pictures showing cells at various stages (D) and cell types generated. (F-I) Immunofluorescence staining of H9 hESC cortical organoids with markers of radial glia (PAX6, BLBP), intermediate progenitor cells (TBR2) and deep layer excitatory projection neurons (TBR1, CTIP2). (j) Spearman’s rank correlation plot of the top 250 upregulated and downregulated genes (H9^NOTCH2NLΔ^ / H9*), and the matching data in W3, W4 and W5 H9 organoids. Numbers in plot indicate pairwise correlation values. (K) Heatmap showing expression profiles for a selection of genes in the significantly enriched GO cluster ‘neuron differentiation’. n = 3.

Neuronal tissues were generated from these clones by directed differentiation of human H9 hESC into cortical organoids (**Figure 5C-D**). hESC-derived cortical organoids resemble early developmental stages of primate cortex development (Eiraku et al., 2008) (**Figure 5E-I**), displaying neural rosette structures that contain radially organized cortical RG cells giving rise to cortical neurons (**Figure 5E-F**) (Lancaster et al., 2013), (Pasca et al., 2015). Bulk RNA-seq transcriptome analysis of cortical organoids isolated at weekly time points reveals that human ESC-derived cortical organoids display efficient and selective induction of dorsal forebrain marker genes, highly resembling the expression pattern during early stages (8-9 post conception weeks) of human dorsal forebrain development *in vivo* (**Figure S5B, Table S2**). H9*^NOTCH2NLΔ^* organoids remained slightly smaller compared to control (H9*) organoids (**Figure S5C**). At week 4 (w4), cortical organoids were harvested for RNA-seq analysis and differentially expressed (DE) genes between H9*^NOTCH2NLΔ^* and H9* were discovered by DESeq2 analysis (Love et al., 2014). To investigate a potential shift in timing of cortical neuron differentiation between H9*^NOTCH2NLΔ^* and H9* organoids, gene expression of the top 250 up- and down-regulated DE genes was correlated to the previously generated RNA-seq profiles of H9 w3, w4 and w5 cortical organoids (Figure S5B, Table S2). This analysis reveals that while differentially expressed genes in w4 H9* correlate best with w4 H9 cortical organoids, strikingly, w4 H9*^NOTCH2NLΔ^* showed a better correlation with w5 H9 cortical organoids (**Figure 5J**). This indicates that w4 H9*^NOTCH2NLΔ^* organoids prematurely express characteristics of older w5 organoids, suggesting H9*^NOTCH2NLΔ^* are advanced in their development compared to H9* organoids. To obtain better insights into what aspects of cortical organoid development were advanced by NOTCH2NL deficiency, we performed GO-term enrichment analysis for the selection of DE genes in H9*^NOTCH2NLΔ^*which were found to correlate better with w5 H9 cortical organoids than organoids of w4 (**Figure 5K**). This cluster of 212 DE genes was found to be enriched for genes involved in neuron differentiation, including key regulators of neuron differentiation such as *DLX1, BCL11B, SEMA3A, UNC5D* and *FGFR2*. These data suggest that w4 H9*^NOTCH2NLΔ^* display premature differentiation, which further supports a potential role for NOTCH2NL in delaying differentiation of neuronal progenitors in the human cortex.

### NOTCH2NL amplifies Notch-signaling through direct interaction with NOTCH receptors

We next sought to test whether *NOTCH2NL* can influence NOTCH2 signaling. The 6 N-terminal EGF domains encoded in NOTCH2NL are also present in the full-length NOTCH2 receptor. These specific domains do not have a clearly described function in activation of the NOTCH pathway. However, they are conserved from *Drosophila* to human, indicating that they are important for normal NOTCH receptor function. There is evidence that the N-terminal EGF domains are involved in dimerization of NOTCH receptors, and that receptor dimerization modulates NOTCH activity (Duering et al., 2011), (Kopan and Ilagan, 2009), (Xu et al., 2015), (Nichols et al., 2007). For example, mutations in the N-terminal EGF domains of NOTCH3 are commonly found in patients suffering from a disease called CADASIL, a hereditary stroke disorder, (Joutel et al., 1997) and these mutations cause aberrant aggregation and dysfunction of NOTCH3 receptors (Karlstrom et al., 2002).

To establish a functional relationship between NOTCH2NL and NOTCH2, we searched for interactions between these proteins. By co-transfecting HA-tagged versions of *NOTCH2NL^Sh^* and *NOTCH2NL^L,T1971^* with a myc-tagged version of NOTCH2 in HEK293 cells we were able to detect co-immunoprecipitation of both proteins by pull-down of either NOTCH2NL or NOTCH2 (**Figure 6A**).

**Figure 6.**
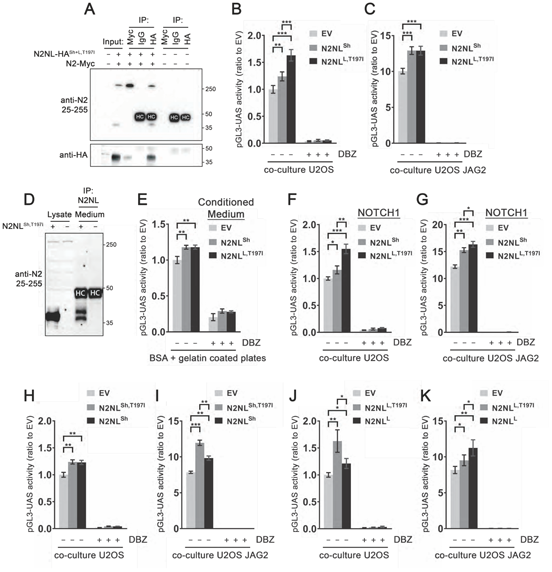
NOTCH2NL paratypes interact with NOTCH receptors and amplify NOTCH signaling. (A) Co-immunoprecipitation of NOTCH2 and NOTCH2NL analyzed by immunoblot. (B-C) Co-transfection of NOTCH2-GAL4 and NOTCH2NL boosts activity of the pGL3-UAS reporter. Upon stimulation of NOTCH2-GAL4 with JAG2 in a coculture setup, NOTCH2NL remains effective in boosting reporter activity. Average of 4 independent experiments with 6 replicates each. Two-way ANOVA with Tukey’s HSD (** p <10^-8^, *** p <10^-12^), error bars indicate SEM. (D) NOTCH2NL is secreted and can be immunoprecipitated from the medium of mouse ESCs ectopically expressing NOTCH2NL. (E) Addition of NOTCH2NL-conditioned medium to cells transfected with NOTCH2-GAL4 and the pGL3-UAS reporter also enhances reporter activity. Average of 2 independent experiments with 4 and 3 replicates each. One-way ANOVA with Tukey’s HSD (* p < 10^-4^), error bars indicate SEM. (F-G) The effect of NOTCH2NL is not restricted to NOTCH2, in an assay with NOTCH1-GAL4 we observe similar effects. (H-K) The presence or absence of the ancestral start codon, and the amino acid 197 Thr™»lie variant, each show specific characteristics in the reporter assays. (H-I) Average of 2 independent experiments with 6 replicates each. Two-way ANOVA with Tukey’s HSD (** p < 10^-8^, *** p < 10^-12^), error bars indicate SEM. (J-K) 6 replicates in one experiment. Student’s t-test with Holm-Bonferroni correction (* p < 0.05, ** p < 10^-3^), error bars indicate SD. See also Figure S6.

Under these co-IP conditions, we did not find detectable interactions with the other EGF-like domain interacting proteins EGFR and PDGFRB (**Figure S6A-B**). These findings indicate NOTCH2NL may influence the conformation and dimerization of NOTCH2.

To assess the influence of NOTCH2NL on activation of the NOTCH2 receptor, a luciferase reporter assay was used. The NOTCH2-ICD DNA binding domain was replaced with a GAL4 domain (NOTCH2-GAL4), and used together with a pGL3-UAS luciferase reporter. CMV-Renilla was used in all reporter experiments for normalization.

This allows for precise measurements of NOTCH2-GAL4 activation, without secondary effects from other NOTCH receptors or other pathways (Habets et al., 2015), (Groot et al., 2014). Co-transfection of empty vector (EV), *NOTCH2NL^Sh^* or *NOTCH2NL^L,T1971^* with *NOTCH2-GAL4* in U2OS cells increased pGL3-UAS reporter activity (**Figure 6B**). *NOTCH2NL^Sh^* increased activity by 24% and *NOTCH2NL^L,T1971^* increased activity by 63%, indicating both the short and long forms of NOTCH2NL can enhance NOTCH receptor activation.

Next, we tested whether NOTCH2NL is still functional under active NOTCH signaling conditions. For this, U2OS cells were transfected with *NOTCH2-GAL4* and either EV, *NOTCH2NL^Sh^* or *NOTCH2NL^L,T1971^.* The transfected cells were then co-cultured on a layer of cells expressing the NOTCH ligand JAG (U2OS-JAG2-Myc) or control (U2OS) cells. Trans-interaction of NOTCH2-GAL4 with JAG2 leads to high activation of the pGL3-UAS reporter in this system. NOTCH2NL was able to increase cleavage of the NOTCH2-GAL4 receptor, indicating that even under these high-signaling conditions NOTCH2NL can further amplify Notch signaling (**Figure 6C**).

We then tested interactions specific to secreted NOTCH2NL. To make secreted NOTCH2NL, mouse ESCs were transfected with *NOTCH2NL^Sh^* or EV as a control. The medium was collected after 32 hours, and used for immunoprecipitation with NOTCH2 aa 25-255 antibody. The isolated protein samples were analyzed by Western blot, which confirmed the presence of secreted NOTCH2NL in the medium (**Figure 6D**). The two bands of *NOTCH2NL^Sh^* may represent the glycosylated protein (higher band) and unmodified protein (lower band). This pattern was also observed in ectopic expression of N-terminal fragments of the NOTCH3 receptor (Duering et al., 2011).

To determine if NOTCH2NL can act cell non-autonomously on NOTCH signaling, we generated medium conditioned with secreted NOTCH2NL or EV-transfected control cells. U2OS cells were transfected with only NOTCH2-GAL4 and the luciferase reporter. After 6 hours, cells were replated in either EV-, *NOTCH2NL^Sh^*- or *NOTCH2NL^L,T1971^*- conditioned medium harvested from other transfected U2OS cells. After 24 hours, cells were isolated for luciferase measurements. NOTCH2NL^Sh^-conditioned medium increased pGL3-UAS reporter activity by 24% and NOTCH2NL^L,T197I^-conditioned medium increased reporter activity by 22% (**Figure 6E**). NOTCH2NL^Sh^ and NOTCH2NL^L,T1971^ have a different N-terminal structure (**Figure 1C**), whereby NOTCH2NL^Sh^ lacks the NOTCH2 signal peptide for canonical transport through the secretory pathway, yet NOTCH2NL^Sh^ is equally potent as NOTCH2NL^l,T1971^ when present in conditioned medium. This suggests NOTCH2NL^Sh^ acts in a cell non-autonomous manner on Notch signaling after being secreted by an unconventional pathway (Steringer et al., 2015), (Rabouille, 2017). In contrast, in the co-transfection experiments NOTCH2NL^l,T1971^ was clearly more potent than NOTCH2NL^l,T1971^-conditioned medium (**Figure 6B**). This suggests NOTCH2NL^L,T1971^ can amplify Notch signaling through both cell-non-autonomous and cell-autonomous mechanisms.

We next tested if NOTCH2NL could also influence the activity of the other NOTCH receptor paralogs NOTCH1 and NOTCH3. NOTCH2NL^Sh^ and NOTCH2NL^L,T1971^ were able to activate cleavage of both NOTCH1-GAL4 and NOTCH3-GAL4, both in coculture with U2OS control cells and U2OS-JAG2 cells, at a level similar to NOTCH2-GAL4 (**Figure 6F-G, Figure S6C**). To assess whether NOTCH2NL functions differently under various ligand stimulated conditions, we tested the ability of NOTCH2NL to enhance NOTCH receptor activity in OP9 cells stably expressing DLL1 and control OP9 cells. NOTCH2NL^Sh^ and NOTCH2NL^lT1971^ mediated activation of the NOTCH2-GAL4 receptor was essentially the same when stimulated by JAG2 or DLL1 (**Figure S6D**). Lastly, NOTCH2NL was similarly able to activate NOTCH2-GAL4 in U2OS cells seeded on recombinant DLL4 coated plates (**Figure S6E**). These data show that NOTCH2NL’s amplifying effect on Notch signaling is not restricted to NOTCH2, and that the potency of NOTCH2NL to enhance NOTCH receptors is not specific for a particular NOTCH-ligand.

Unlike NOTCH2NL^Sh^, NOTCH2NL^L,T1971^ harbors a common variant present in EGF domain 5 that changes the ancestral threonine at position 197 to isoleucine (**Figure 1C**). This threonine can be fucosylated, while isoleucine cannot. Fucosylation is essential for modifying binding of EGF domains to their interaction sites. We tested the short and long versions of NOTCH2NL with and without the T197I variant in the NOTCH2-GAL4 - pGL3-UAS reporter assay. Interestingly, the single amino acid changes had subtle but significant influences on NOTCH2-GAL4 cleavage. *NOTCH2NL^Sh^* and *NOTCH2NL^Sh,T1971^* performed equally under baseline conditions (29% and 31% increase, respectively, **Figure 6H**). However, with JAG2 co-culture, *NOTCH2NL^Sh,T1971^* variant was more potent than *NOTCH2NL^Sh^* (53% and 25% increase respectively, **Figure 6I**). *NOTCH2NL^L,T1971^* was also a more potent activator than *NOTCH2NL^L^* (63% and 21% increase, respectively, **Figure 6J**). However, co-culture with JAG2 cells showed the opposite. Here, *NOTCH2NL^L,T1971^* had less effect than *NOTCH2NL^L^* (17% and 38% increase, respectively, **Figure 6K**).

Altogether, these data suggest that NOTCH2NL can enhance NOTCH receptor activation. Structural changes between the various NOTCH2NL paratypes, varying from the absence/presence of the signal peptide to single amino acid changes in the EGF-like repeats, show differential effects under different conditions. This implies that the influence of NOTCH2NL genes on cortical neurogenesis in any particular individual may be dependent on the combination of NOTCH2NL paratypes he or she carries. However, widespread genomic variation in NOTCH2NL paratypes in the healthy human population shows that the absence or loss of one specific paratype is not necessarily associated with disease, so there must be some robustness to the system as well.

### Deletions and duplications of NOTCH2NL genes are associated with neurodevelopmental phenotypes

The revision of the sequence of the 1q21.1 band in the human reference genome GRCh38 for the first time clearly separated the Thrombocytopenia Absent Radius (TAR) syndrome’s centromerically proximal locus from the 1q21.1 deletion/duplication syndrome’s distal locus (Figure 1A). The latter is bracketed by *NOTCH2NLA* at GRCh38.chr1:146,151,907-146,229,032 (within a remapped region that was previously called breakpoint 3) and *NOTCH2NLB* at GRCh38.chr1:148,602,849-148,679,774 (aka breakpoint 4) (Mefford et al., 2008), (Rosenfeld et al., 2012) (**Figure S7**).

These breakpoints are 99.7% identical over an interval of approximately 250,000 bases, creating frequent non-allelic homologous recombination that results in either deletion or duplication of a 2.4 MB segment in the chromosome in which it occurs. Supporting the high frequency of the event, it is estimated that 18-50% of distal duplications and deletions seen in the clinic are *de novo* (Haldeman-Englert and Jewett, 1993). The remainder inherit the condition, often from an asymptomatic or mildly symptomatic parent. The frequency of the deletion in the European general population has been estimated at 0.03% and the duplication at 0.049% (Mace et al., 2017).

When present, symptoms are strikingly, but not exclusively, neurological. In several independent large studies 1q21.1 distal deletion has been shown to be associated with schizophrenia (International Schizophrenia Consortium, 2008; Mace et al., 2017), (Grozeva et al., 2012), (Chang et al., 2016). In contrast, in patients ascertained as part of the Simons Variation in Individuals (VIP) autism study, 1q21.1 distal duplication probands exhibited ADHD (29%), behavior disorder (18%), autism spectrum disorder (41%), developmental coordination disorder (23%), intellectual disability (29%), while deletion probands exhibited these symptoms at lower frequencies, but with a relatively high percentage (26%) exhibiting anxiety and mood disorders (Bernier et al., 2016). Of particular interest regarding the function of NOTCH2NL in the enhancement of NOTCH signaling during neurodevelopment is the fact that studies have consistently shown that the duplication is associated with macrocephaly and the deletion with microcephaly (Mefford et al., 2008), (Brunetti-Pierri et al., 2008).

Because the copy number variation studies that identified the 1q21.1 syndromes were performed on microarrays that were based on the GRCh37 and earlier assemblies that were incorrect in the 1q21.1 band, we reanalyzed copy number variation microarray data derived from 11 previously characterized patients where remappable data could be obtained, 9 with microcephaly and 2 with macrocephaly, to determine the boundaries of their 1q21.1 variations in GRCh38 coordinates. Probe positions were realigned to the revised assembly, and a more sensitive method of copy number variation analysis involving optimization by integer linear programming was developed (STAR Methods). The remapped data suggest that *NOTCH2NLA* and *NOTCH2NLB* are located inside the 1q21.1 locus implicated in microcephaly and macrocephaly (**Figure S7**). Nine out of the nine microcephalies were consistent with *NOTCH2NLA* and/or -*B* deletion and both macrocephalies were consistent with *NOTCH2NLA* and/or *-B* duplication. In at least one out of nine microcephaly patients (number 2), the *HYDIN2* locus exhibits a normal copy number, consistent with results previously published (Dougherty et al., 2017).

The sparsity of unique probes in the highly repetitive 1q21.1 region precludes a precise breakpoint analysis of these older data. In order to better map the breakpoints of the copy number variations in 1 q21.1 patients, we obtained primary fibroblasts from six autism patients in the Simons VIP project (Bernier et al., 2016) one listed as having 1q21.1 distal duplication and five identified as having the distal deletion (**Figure 7, Table S3**), and prepared high molecular weight DNA for targeted sequencing and assembly as described above. The sequencing coverage data demonstrates that all of these samples have their deletion or duplication in the breakpoint 3 and 4 regions containing *NOTCH2NLA* and *NOTCH2NLB* (**Figure 7A**). In particular, in deletions we see normal copy number 2 for unique DNA outside of the locus and copy number 1 for unique DNA inside the locus. In the two near-identical breakpoint regions that flank the locus we expect both NOTCH2NLA and NOTCH2NLB to be present on the non-deleted chromosome, and a homology-driven deletion in the other chromosome is expected to leave a single NOTCH2NL A/B gene, possibly a fusion gene (**Figure 7B**). Indeed, we see combined read coverage 3 for both breakpoint regions, which is split into two, leaving an “intermediate level plateau” of read depth at roughly 1.5 in each of the NOTCH2NL regions. In the case of a duplication, these intermediate level plateaus at the breakpoint flanks are at read depth 2.5, implying 5 copies of *NOTCH2NLA* and *B*.

**Figure 7.**
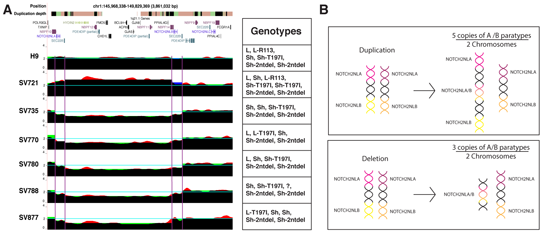
Patients with 1q21.1 Distal Deletion/Duplication Syndrome show breakpoints and CNV in *NOTCH2NLA* and *NOTCH2NLB*. (A) UCSC Genome Browser screenshot from GRCh38. The duplication depth track indicates duplicated genome sequences as colored bars: white (single copy, N=1), orange (N=2-4), green (N=5), black (N > 5). The normalized read depth for each individual is in red (see methods). For breakpoints in *NOTCH2NL,* due to sequence similarity and gene conversion, we expect the average normalized coverage of *NOTCH2NLA* and *NOTCH2NLB* to be 1.5 for deletions and 2.5 for duplications. Similarly, we expect the normalized coverage of the sequence in between to be 1.0 for deletions and 3.0 for duplications (blue). The region was then broken into 5 portions -- centromeric to *NOTCH2NLA, NOTCH2NLA,* in between *NOTCH2NLA* and *NOTCH2NLB, NOTCH2NLB* and telomeric to *NOTCH2NLB.* The average values of the normalized coverage based on the five regions are drawn in green. Where all models agree, the colors combine to black. (B) Schematic of the *NOTCH2NL* chromosomal configuration before and after a duplication or deletion event. In the case of duplication, there is an extra NOTCH2NL paratype that is a hybrid of the *NOTCH2NL* paratype at the *NOTCH2NLA* position and that at the *NOTCH2NLB* position from that position. For a deletion, only a hybrid of the two *NOTCH2NL* paratypes from that chromosome remains. See also Figure S7, Table S3-S4.

When we assembled the *NOTCH2NL* paratypes in these individuals we confirmed that indeed all of the deletion samples have three *NOTCH2NLA* or *NOTCH2NLB* alleles and the duplication sample has five. In all but one sample (SV788) we could determine all of the *NOTCH2NLA/B* paratypes as well (**Figure 7, Table S1**). As expected from the position of the breakpoints in the coverage analysis, all 6 individuals showed no evidence of gain or loss of *NOTCH2NLC*, which has two *NOTCH2NL^Sh-2ntdel^* alleles. The patients, like normal individuals, show a variety of combinations of *NOTCH2NL* paratypes. If there is any pattern in the set of *NOTCH2NL* paratypes that is particular to 1q21.1 distal deletion/duplication syndrome, this patient set is too small to discern it. Nevertheless, this analysis suggests that the typical 1q21.1 distal deletion/duplication syndrome events are driven by non-allelic homologous recombination between *NOTCH2NLA* and *B* or nearby sequence, and these events produce *NOTCH2NL* copy number change. Thus, it remains possible that *NOTCH2NL* copies not only provide the breakpoints for these events, but may have a contribution to the neurodevelopmental phenotypes associated with them as well.

## Discussion

*NOTCH2NL* joins *ARHGAP11B* (Florio et al., 2015) and *BOLA2* (Nuttle et al., 2016) as a third example of the concomitant emergence of a potential adaptive evolutionary innovation and susceptibility to a recurrent genomic disorder from chromosomal instability mediated by hominin-specific duplication hubs. These duplicated regions not only represent the bulk of human-specific genomic DNA (Cheng et al., 2005), but are an important a reservoir of new genes, carrying with them the potential to rapidly change the structure of the genome via non-allelic homologous recombination. Indeed the most promising genes recently implicated in human brain evolution in addition to those above, such as *SRGAP2C* (Dennis et al., 2012), (Charrier et al., 2012) and *TBC1D3* (Ju et al., 2016) map to regions of human-specific structural variation.

The peculiar evolutionary history of *NOTCH2NL* includes a series of complex genomic reorganization events that eventually led to the creation of three functional NOTCH-related genes early in hominin evolution. *NOTCH2NL*-like pseudogenes lacking coding potential, including a *PDE4DIP-NOTCH2NL* pseudogene, were already present in the gorilla, chimpanzee and human LCA and still remain in the genomes of chimpanzee and gorilla, but are not found in hominin genomes. Instead, in each of the three hominin species sequenced, human, Neanderthal and Denisovan, we find 3 *NOTCH2NL* genes, *NOTCH2NLA, B* and *C*, and in humans we find an additional pseudogene, *NOTCH2NLR.* The most plausible explanation for this is that in a common ancestor of humans, Neanderthals and Denisovans, the ancestral *PDE4DIP-NOTCH2NL* pseudogene was repaired by ectopic gene conversion from *NOTCH2.* This event may have been crucial to hominin evolution, as it marks the birth of a novel hominin-specific NOTCH-related gene that likely functions to delay yet eventually increase production of neurons from neural stem cells during fetal brain development, and whose introduction was prior to or during a time of rapid expansion of brain size in hominins approximately 3 MYA (Halloway, et al., 2004) (**Fig. S2E**). Subsequent duplication events created additional *NOTCH2NL* genes in hominin ancestors, increasing *NOTCH2NL* dosage. In modern humans, three *NOTCH2NL* genes are present with at least eight different “paratypes”, i.e., haplotype configurations that can occur in different paralogous locations.

Consistent with a potential role in brain development, *NOTCH2NL* is expressed in the germinal zones of the developing human cortex where it likely promotes NOTCH signaling. Of particular interest is the expression of NOTCH2NL in outer radial glia cells, which are responsible for a more pronounced proliferative zone in the human developing cortex compared to that in related primates and are thought to be involved in the evolutionary expansion of the human cortex (Hansen et al., 2010, Lui et al., 2011). It has remained elusive how these loosely organized oRG cells maintain their sustained capacity for self-renewal and proliferation, as they lie in the outer subventricular zone (OSVZ) far away from the proliferative zone of strictly laminar radial glial cells near the surface of the ventricle. Recent work suggests that oRG cells directly contribute to a stem cell niche of the OSVZ through increased expression of extracellular matrix proteins and growth signal factors (Pollen et al., 2015). It has been shown that Notch signaling is essential for oRG cell self-renewal (Hansen et al., 2010), and our data suggest NOTCH2NL may act as a potentiating factor on Notch signaling to support oRG cell self-renewal. In particular, we show that NOTCH2NL can act in part in a cell non-autonomous manner to amplify Notch signaling. Our data also supports a separate, cell-autonomous function of NOTCH2NL, and indeed, in a very recent related study (Suzuki, et al, Submitted), it was found that the long version of NOTCH2NL (NOTCH2NL^L,T197L^) promotes NOTCH signaling and neural progenitor proliferation in a cell-autonomous manner. Through these actions NOTCH2NL could support self-renewal of oRG cells in the OSVZ environment.

Enhancement of NOTCH signaling is likely the mechanism that underlies the gene expression phenotypes we observed in experiments in which NOTCH2NL is ectopically expressed in mouse or is deleted in human cortical organoids. We showed that stable expression of NOTCH2NL in mouse organoids leads to downregulation of genes involved in neuronal differentiation and upregulation of genes involved in negative regulation of neuronal progenitor differentiation. These results are further supported in human cortical organoids grown from hESC lines with five out of six alleles of *NOTCH2NLA/B/C* deleted by CRISPR/Cas9, where NOTCH2NL deficiency resulted in premature expression of genes involved in neuronal differentiation.

The combined role of NOTCH2NL genes in any particular genotype may depend on the combined expression pattern of structurally distinct NOTCH2NL paratypes. The short forms of NOTCH2NL may act primarily in a cell non-autonomous way, as they are likely secreted by unconventional secretion pathways due to the lack of a signal peptide. Several routes of unconventional protein secretion have been described (Rabouille, 2017), of which FGF2 secretion is a well-studied example (Steringer et al., 2015). In contrast, the long forms of NOTCH2NL contain the signal peptide required to enter the canonical secretory pathway, where they can encounter NOTCH receptors and NOTCH ligands in secretory vesicles before they are transported to the outer membrane of the cell. The long forms of NOTCH2NL therefore may act primarily cell autonomously or may act both autonomously and non-autonomously. We found that the different paratypes of NOTCH2NL have different potencies to amplify NOTCH activity and do so under different circumstances (presence or absence of Notch ligand, secreted in the medium or generated within the cell, etc.). This suggests there may have been an evolutionary gene-dosage optimization, where NOTCH2NL paratypes with different structural and sequence properties have undergone multiple changes to reach a balanced state of NOTCH pathway modulation.

The existence of many distinctly functional NOTCH2NL paratypes, created by substitutions and small indels that were likely to have been combinatorially amplified by ectopic gene conversion during hominin evolution, is striking. While this has resulted in many versions of NOTCH2NL encoded at the *NOTCH2NLA* and *NOTCH2NLB* positions in chromosome 1q21, the overall copy number of *NOTCH2NLA and NOTCH2NLB* is remarkably stable at a combined copy number 4 in almost all the 266 individuals of the Simons Diversity collection, with some being ambiguous. This suggests total dosage of these A/B paratypes may be important. In contrast, *NOTCH2NLC* has distinct paratypes (characterized primarily by a 2 bp deletion that forces use of a downstream start codon, denoted *NOTCH2NL^Sh-2ntdel^*), and does not show as stable a copy number in the Simons collection (appearing to have copy number 3 in 3 individuals, copy number 1 in 20 individuals, and 0 in 1 individual). Further, despite the apparent conservation of A/B copy number, one of the A/B paratypes, *NOTCH2NL^LR113*^*, contains a premature stop codon that is predicted to render the protein nonfunctional. This paratype appears to be present at copy number 2 in 4 individuals and copy number 1 in 31 in the Simons collection. There also appear to be 4 individuals who have 1 copy of *NOTCH2NL^LR113*^* and lack one copy of *NOTCH2NL^Sh-2ntdel^,* so copies of *NOTCH2NL^Sh-2ntdel^* in the C paralog position are not always there to make up for stop codon-containing copies in the A or B positions.

Detailed modeling of selective forces on NOTCH2NL is complicated by the relatively small number of individuals whose genotypes have been resolved into a specific combination of paratypes, and the peculiar mix of substitution and ectopic gene conversion between equivalent loci that creates a diversity of paratypes. Lacking large scale sequencing with long DNA fragments, the selective forces on *NOTCH2NL* genotypes remain mysterious. Still, it is clear that *NOTCH2NL* genotypes exist with different dosages of different functionally distinct paratypes in the population. Either the physiological system is quite robust to these protein differences despite their differential effect on Notch signaling, or the optimal balance of *NOTCH2NL* alleles has not been fixed in the population.

Despite their presence in the breakpoints of the 1q21.1 distal deletion/duplication region, *NOTCH2NL* genes have not previously been associated with this syndrome because the reference genome assembly was incorrect until the GRCh38 assembly. By reanalyzing older data and generating new data from patients from whom long genomic DNA fragments could be isolated, we find that all patients we have examined have copy number variations that include an extra *NOTCH2NL* copy in the duplication and loss of a copy in the deletion. Further, wherever it was possible to determine, we find that non-allelic homologous recombination has occurred inside or near *NOTCH2NLA* and B, leaving behind an intact *NOTCH2NL* copy, possibly a chimera of A and B (**Figure 7B**). Therefore, *NOTCH2NLA* and *B* should be considered candidates for contributing to the phenotypes of 1q21.1 deletion/duplication syndrome, including abnormal brain size phenotypes (Mefford et al., 2008), (Brunetti-Pierri et al., 2008), (Dumas et al., 2012), (Girirajan et al., 2013). However, they are not the only genes whose copy number is changed. Protein coding genes between *NOTCH2NLA* and *NOTCH2NLB* include *HYDIN2, CHD1L, BCL9, GJA5, GJA8, PRKAB2, FMO5, ACP6, PPIAL4G, GPR89B*, and DUF1220-domain containing *NBPF11, NBPF12 and NBPF14* paralogs, three of the more than fourteen functional paralogs of *NBPF*. The progressive increase in number of DUF1220 domains in primate genomes has been shown to correlate with the evolutionary expansion of the neocortex (Popesco et al., 2006), and *CHD1L* is amplified in many solid tumors and promotes tumor growth (Cheng et al., 2013), (Xu et al., 2013), thus it could be associated with cell proliferation. *HYDIN2* was the strongest candidate for brain size effects of 1 q21.1 distal deletion/duplication syndrome as it shares a protein domain with the microcephaly-associated protein *ASPM* and its paralog *HYDIN* lies in the 16q22.2 locus that has also been associated with microcephaly. However, recent work found 6 symptomatic patients with atypical breakpoints excluding *HYDIN2*, thus eliminating it as a likely driver (Dougherty et al., 2017). Mapping of these atypical patients indicates that the *NOTCH2NLA* locus was also excluded in these atypical events, but the *NOTCH2NLB* locus was fully duplicated in the three atypical duplication cases (all macrocephalic), and fully deleted in the three atypical deletion cases (all microcephalic). Therefore, these cases are consistent with a possible contribution of NOTCH2NL to 1 q21.1 deletion/duplication syndrome. There are scattered reports of patients with smaller duplications or deletions, a few of these including neither *NOTCH2NLA* nor *NOTCH2NLB* (Girirajan et al., 2013), (Van Dijck et al., 2015). These studies have variously pointed to *CHD1L, BCL9* or the noncoding RNA gene *LINC00624* as candidates for causing 1q21.1 deletion/duplication syndrome symptoms. However, no consistent pattern has been found and no mechanistic explanation for the possible association of these genes with neurodevelopmental changes has yet been established. 2.4 megabases is a considerable stretch of DNA, and it may well be that these or other genes, such as the three *NBPF* paralogs in the region, contribute to the neurological effects of this large copy number change. Indeed non-neurological 1q21.1 distal deletion/duplication effects in the heart and eyes are thought to be caused by the copy number changes in *GJA5* and *GJA8* (Gollob et al., 2006) (Shiels et al., 1998), so it is plausible that several genes may contribute to the apparently complex neurological effects. At the very least, by providing the substrates for non-allelic homologous recombination, *NOTCH2NL* genes enable the 1 q21.1 distal deletion/duplication syndrome.

The strongly directional association of 1q21.1 distal deletion/duplication syndrome with brain size, with duplications tending to cause macrocephaly and deletions microcephaly, is most intriguing from a NOTCH2NL perspective, given that duplications are associated with increased NOTCH2NL dosage, which in *ex vivo* experiments delays differentiation of neuronal progenitor cells allowing for a longer period of proliferation, and deletions decrease NOTCH2NL dosage, which promotes premature differentiation of neuronal progenitor cells. While still circumstantial, these data support a role of *NOTCH2NL* genes not only in the cognitive and brain size symptoms of 1 q21.1 distal deletion/duplication syndrome, but in the evolution of the hominin-specific larger brain size and associated cognitive traits. A delay in cortical maturation, coupled with a net increase in the size of the neocortex, in large part from sustained activity of outer radial glia progenitor cells, is characteristic of human brain development. Humans may in fact be caught in an evolutionary compromise in which having multiple identical copies of *NOTCH2NL* provides a neurodevelopmental function we need while at the same time predisposing our species to recurrent *de novo* non-allelic homologous recombination events that underlie a neurodevelopmental syndrome and contribute to our overall genetic load. Given the many different alleles (“paratypes”) of *NOTCH2NL* we observe segregating in the current human population, the tension caused by this compromise may still be a factor in our ongoing evolution.

## Acknowledgements

We thank Robert Kuhn for help in obtaining data from researchers in the ISCA and DECIPHER consortia, and Mark Diekhans, Brain Raney, and Hiram Clawson for help with features of the UCSC Genome Browser used in this research; Raphael Bernier for help interpreting the Simons VIP phenotype data; Mari Olsen (Haussler lab), Bari Nazario (the Institute for the Biology of Stem Cells), Nader Pourmand (the UCSC Genome Sequencing Center), Ben Abrams (the UCSC Life Science Microscopy Center) and Shana McDevitt (UC Berkeley, QB3, Vincent J. Coates Genomics Sequencing Laboratory) for their excellent technical support; Arjan Groot and Marc Vooijs for generous supply of reagents and support for reporter assays; Martijs Jonker (SILS, University of Amsterdam) for help in RNA-seq data processing; Pierre Vanderhaeghen, Bin Chen, Benedict Paten, Ed Green and members of the Haussler and Jacobs labs provided helpful discussions and comments on the manuscript. This work was supported by CIRM Predoctoral Fellowship T3-00006 (ARF), CIRM Postdoctoral Fellowship TG2-01157 (FMJJ), Human Frontier Science Program Postdoctoral Fellowship LT000689/2010-L and HFSP Career Development Award CDA00030/2016C (FMJJ), Marie Curie reintegration grant H2020-MSCA-IF-2014_RI 659193 (FMJJ), NWO Earth and Life Sciences (ALW), project 834.12.003., EMBO ALTF 292-2011 (MH) and a special fellowship from donor Edward Schulak (ADE), NIH HG002385 (EEE), NIH F30HG009478 (MLD), CIRM Center of Excellence for Stem Cell Genomics (Stanford) GC1R-06673-A, CIRM Center for Big Data in Translational Genomics (SALK) GC1R-06673-B, and StemPath NIH/NIGMS R01 GM109031 (DH), NCI Cloud award fund #24074-443720 (DH), and the California Institute for Quantitative Biosciences. David Haussler and Evan E Eichler are Investigators of the Howard Hughes Medical Institute.

## Author Contributions

Conceptualization- D.H., S.R.S., F.M.J.J., I.T.F.; Methodology- I.T.F., S.R.S., D.H., M.M., A.D.E., G.L.M., A.Bi., A.M.N., F.M.J.J.; Validation- M.J.D., X.N.; Investigation-G.A.L., M.M., G.L.M., Avd.B., J.L.R., A.R.F., L.R., T.J.N., A.A.P.; Formal Analysis- I.T.F., A.D.E., A.M.N., A.B., R.L.R., A.B., M.L.D., X.N., S.K.; Resources-T.J.N., A.A.P., M.C.A., S.Z., E.E.E., A.K.; Data Curation- I.T.F., C.M.B., G.A.L.; Writing-Original Draft- D. J., I.T.F., G.A.L., F.M.J.J., S.R.S.; Writing-Review & Edit- D.J., I.T.F., G.A.L., F.M.J.J., S.R.S., C.M.B., A.D.E., G.L.M., A.R.F., M.H., T.J.N., A.A.P., M.L.D., X.N.; Visualization- I. T.F., G.A.L., G.L.M., A.M.N., M.H.; Supervision- A.K., E.E.E., S.R.S., F.M.J.J., D.H.; Project Administration- D.H., F.M.J.J.; Funding acquisition- D.H., F.M.J., E.E.E.

## Supplemental Figure Legends (See “Supplemental Materials.pdf” for the figures)

**Figure S1. Related to Figure 1**. (A) Identity between NOTCH2-related genes as measured over the alignable genomic region (blue) or the gene exons (pink). (B) DNA and amino acid sequence of human NOTCH2NL genes in exon 5, which is derived from *NOTCH2* intron 4. (C) Immunoblot of mouse ESCs transfected with WT *NOTCH2NL,* or *NOTCH2NL* with ancestral ATAA inserted in exon 5. (D) Relative protein levels based on the immunoblot in panel C. (E) RT-qPCR analysis of the same samples for determination of transcript levels for each condition. (F) Alignment of the H9 assembled paratypes to GRCh38. Each paratype is colored as to whether a position aligns best to GRCH38 *NOTCH2NLA, NOTCH2NLB* or *NOTCH2NLC.* (G) Observed frequency of individuals with the indicated *NOTCH2NLC* and *NOTCH2NLR* copy number in the Simons Diversity Panel (n=266). (H) Schematic of linked-read sequencing and Gordian Assembler protocol using the 10X genomics Chromium genome assay and oligocapture to enrich for library fragments containing the desired genomic region. (I) Protein alignment of observed NOTCH2, NOTCH2NL and NOTCH2NLR paratypes based on our assembly results. Note that NOTCH2 sequence extends beyond what is shown in the alignment. A segregating variant in NOTCH2NLR is found at amino acid position 235.

**Figure S2. Related to Figure 2**. Details of *NOTCH2NL*-like genes in Gorilla (A) and Chimp (B) and source of genome sequence support. (C) Verification of fusion genes by RT-PCR on Chimp and Gorilla RNA. H = Human, C = Chimpanzee, G = Gorilla. (D) Immunoblot blot using an N-terminal NOTCH2 antibody (aa 25-255), comparing ectopic expression of Chimp *NOTCH2NL*-like gene lacking exon 2 compared to ectopic expression of human *NOTCH2NL^Sh,T197I^* in mESCs. (E) Endocranial volume of fossil hominids versus time as determined by Holloway, et al. 2004. (F) Details of splice junctions of fusion genes and related open reading frames. Top rows show nucleotide sequence in fusion transcripts. Middle rows show peptides derived from these transcripts. Lower rows contain the orthologous human NOTCH2NL protein sequence.

**Figure S3. Related to Figure 3**. (A) Summary violin plots indicating *NOTCH2NL* and *NOTCH2* expression in various cell types. (B) *NOTCH2NL* paratype expression in undifferentiated hESCs and week 5 cortical organoids from bulk Illumina RNA-Seq.

**Figure S4. Related to Figure 4**. (A) MA plot of RNA-sequencing data of mouse cortical organoids based on DESeq2 analysis. (B) Z-score of differentially expressed genes (p-adj < 0.05, DESeq2). (C) GO terms significantly associated with the upregulated genes in organoids ectopically expressing *NOTCH2NL^Sh,T1971^*.

**Figure S5. Related to Figure 5**. (A) Schematic of strategy to generate *NOTCH2NL-* specific deletions using CRISPR/Cas9 and alignment of the two guide sequences used to NOTCH2NL-related sequences. (B) Heatmap of expression levels for a selection of brain structure marker genes from hESC-derived cortical organoids at the indicated time points (Left) and from human embryonic dorsal prefrontal cortex (DFC) samples at 8 pcw, 9 pcw, 12 pcw and 13 pcw, derived from the Allen Brain Atlas (http://www.brainspan.org). w = week; pcw = post conception week; FPKM= fragments per kilobase of exon per million fragments mapped. (C) Brightfield images of developing H9* and H9^NOTCH2NLΔ^ cortical organoids. Day 28 images show organoid pools for 1, RNA-Seq replicate.

**Figure S6. Related to Figure 6**. (A-B) Investigation of co-immunoprecipitation of NOTCH2NL with PDGFRB and EGFR in two independent experiments. (C) Reporter assay to assess the effect of NOTCH2NL using either NOTCH1-GAL4, NOTCH2-GAL4 or NOTCH3-GAL4 to induce pGL3-UAS reporter activation. 6 replicates in one experiment. Student’s t-test with Holm-Bonferroni correction (* p < 0.05, ** p < 10^-3^, *** p < 10^-5^), error bars indicate SD. (D) NOTCH2NL effects of NOTCH2-GAL4 reporter assay co-culture with JAG2 or DLL1 expressing cells. 6 replicates in one experiment. Student’s t-test with Holm-Bonferroni correction (* p<0.05, ** p<10^-3^, *** p<10^-5^), error bars indicate SD. (E) NOTCH2NL remains effective in an assay with activation of NOTCH2-GAL4 using recombinant DLL4 coated plates. Average of 3 independent experiments with 4 or 5 replicates each. Two-way anova with Tukey’s HSD (* p < 10^-4^, ** p <10^-8^, *** p < 10^-12^), error bars indicate SEM.

**Figure S7. Related to Figure 7**. Relative probe intensities from CNV-microarrays for 2 controls and 11 patients with reported 1q21.1 aberrations mapped to the GRCh38 1q21.1 assembly. Gray: normal, red: deletion, blue: duplication. Dark red/blue is high confidence deletion/duplication based on probe values, light red/blue are potentially part of the deletion/duplication.

**Table S1. Related to Figures 1**. Results of NOTCH2NL gene de novo assembly.

**Table S2. Related to Figure 4 and 5**. Gene expression measurements from RNA-seq experiments.

**Table S3. Related to Figure 7**. Features of Simons VIP samples.

**Table S4. Related to Figures 1 and 7**. Curated Paratypes of Assembled and Simons Normals

## STAR Methods

### Mouse stem cell culture and organoid differentiation

Mouse 46c ESCs were grown according to the BayGenomics protocol (Skarnes, 2000), on 0.1% gelatin coated plates, and cultured in GMEM (Thermofisher) with 10% HIFBS, 2 mM L-Glutamine, 1x NEAA, 1x NaPyr, 100 μM 2-mercaptoethanol and 1x P/S. ESGRO LIF (Millipore) was added fresh daily. To generate stable cell lines, 46C cells seeded on 100 mm plates and were transfected with 24 μg of linearized pCIG-NOTCH2NL-ires-GFP or empty pCIG-ires-GFP vector, using lipofectamine 2000 (Thermofisher). After 36 hours, GFP-positive cells were sorted using a FACSAria III (BD Biosciences) and recovered for further culturing. After 4 passages sorting was repeated and GFP-positive cells that had stably integrated the plasmid DNA in their genome were recovered for expansion and further culturing. We verified continued stable expression of NOTCH2NL-ires-GFP or empty vector (Supplemental Figure S4). Mouse 46C ESC organoid differentiation was performed as described previously (Eiraku and Sasai, 2011). Three pools of 16 organoids were isolated in TRIzol after 6 days of differentiation for EV and NOTCH2NL organoids.

### Human stem cell culture

Mitomycin-C treated mouse embryonic fibroblasts (MEFs, GlobalStem) were seeded on 0.1% gelatin coated plates at a density of 35.000 cell/cm^2^. MEFs were cultured in DMEM, 4.5 g/l glucose + GlutaMax (Thermofisher, 10% heat inactivated fetal bovine serum (HIFBS, Thermofisher), 1x Penicillin/Streptomycin (P/S, Thermofisher) and 1x sodium pyruvate (NaPyr, Thermofisher). H9 human embryonic stem cells (H9 hESCs, WiCell), were cultured in W0 medium: DMEM/F12 (Thermofisher) with 20% KnockOut serum replacement (KOSR, Thermofisher), 2 mM L-glutamine (Thermofisher), 1x non-essential amino acids (NEAA, Thermofisher), 100 μM 2-mercaptoethanol (Thermofisher) and 1x P/S (Thermofisher). W0 was freshly supplemented daily with 8 ng/ml FGF2 (Sigma). H9 hESCs were grown on MEF feeder layers, and manually passaged every 5-6 days when colonies reached approximately 2 mm in diameter.

### Human organoid differentiation

For organoid differentiation, medium was replaced with W0 medium + 1x NaPyr without FGF2 (Differentiation medium). Colonies of 2-3 mm in diameter were manually lifted using a cell lifter, and transferred to an ultra-low attachment 60mm dish (Corning). After 24 hours (day 0) embryoid bodies had formed, and 50% of medium was replaced with Differentiation medium supplemented with small molecule inhibitors and recombinant proteins to the following final concentrations: 500 ng/ml DKK1 (peprotech), 500 ng/ml NOGGIN (R&D Systems), 10 μM SB431542 (Sigma) and 1 μM Cyclopamine V. californicum (VWR). Medium was then replaced every other day until harvest. On day 8, organoids were transferred to ultra-low attachment U-shaped bottom 96 well plates (Corning). On day 18, medium was changed to Neurobasal/N2 medium: Neurobasal (Thermofisher), 1x N2 supplement (Thermofisher), 2 mM L-Glutamine, 1x P/S, supplemented with 1 μM Cyclopamine. From day 26 on, Cyclopamine was not supplemented anymore. Organoids were harvested in TRIzol at weekly time points. Total-transcriptome strand-specific RNA sequencing libraries were generated using dUTP for second strand synthesis on Ribo-zero depleted total RNA (Parkhomchuk et al., 2009). For organoid formation of H9 hESC CRISPR/Cas9 NOTCH2NL knockout lines, an updated protocol was used: Differentiation medium was supplemented with 10 μM SB431542 (Sigma), 1 μM Dorsomorphin (Sigma), 3 μM IWR-1-Endo (Sigma) and 1 μM Cyclopamine (Sigma). Medium was then replaced every other day until harvest. On day 4, 60 mm dishes with organoids were placed on a hi/lo rocker in the incubator. From day 18 on, medium is replaced with Neurobasal/N2 medium. From day 24 on, Cyclopamine was not added anymore. Pools of 5-10 organoids per replicate were harvested in TRIzol at day 28 for RNA extraction.

### RNA-Sequencing Analysis

Paired-end Illumina reads were trimmed from the 3’ end of read1 and read2 to 100x100 bp for human. Bowtie2 v2.2.1 (Langmead and Salzberg, 2012) was used with the “--very-sensitive” parameter to filter reads against the RepeatMasker library (http://www.repeatmasker.org) which were removed from further analysis. STAR v2.5.1b (Dobin et al., 2013) was used to map RNA-seq reads to the human reference genome GRCh37. STAR was run with the default parameters with the following exceptions: -- outFilterMismatchNmax 999, --outFilterMismatchNoverLmax 0.04, --alignIntronMin 20, --alignIntronMax 1000000, and --alignMatesGapMax 1000000. STAR alignments were converted to genomic position coverage with the bedtools command genomeCoverageBed -split (Quinlan and Hall, 2010). DESeq2 v1.14.1 (Love et al., 2014)) was used to provide basemean expression values and differential expression analysis across the time course. Total gene coverage for a gene was converted to read counts by dividing the coverage by N+N (100+100) since each paired-end NxN mapped read induces a total coverage of N+N across its genomic positions. Results are in **Table S2** and data are available from GEO: GSE106245.

For mouse cortical organoids, H9* and H9^NOTCH2NLΔ^ organoid samples, RNA was isolated according to standard TRIzol protocol. RNA was treated with DNAseI (Roche) according to standard protocol for DNA clean-up in RNA samples. RNA was then isolated by column purification (Zymo RNA clean & concentrator 5) and stored at −80°C. For RNA sequencing, mRNA was isolated from total RNA using polyA selection Dynabeads mRNA DIRECT Micro Purification Kit (Thermofisher). Library was prepared using strand-specific Ion Total RNA-seq Kit v2 (Thermofisher) and Ion Xpress RNA-seq Barcode 1-16 (Thermofisher) to label different samples. The samples were sequenced on an IonProton system (Thermofisher), generating single-end reads of around 100 bp in length. IonProton RNA sequencing data was processed using the Tuxedo package, in consideration of parameters recommended by Thermofisher for IonProton data. Briefly, samples were mapped using Tophat2 (Kim et al., 2013), using Bowtie2 (Langmead and Salzberg, 2012) as the underlying alignment tool. The target genome assembly for these samples was GRCh38/UCSC hg38, and Tophat was additionally supplied with the gene annotation of ENSEMBL84 (GRCh38.p5). Reads mapped per exon were counted using HT-Seq count (union) and summed per corresponding gene. HT-Seq count output was normalized using DESeq2, and pairwise comparisons were made to determine significant differences in gene expression. Results are in **Table S2**.

For comparison of week 4 H9* and H9^NOTCH2NLΔ^ organoid data to the previously established H9 organoid timeline, the following procedure was used: The top 250 upregulated and the 250 downregulated genes between week 4 H9* and H9^NOTCH2NLΔ^ based on p-adj were selected. The matching expression profiles of these 500 genes were extracted from the H9 organoid timeline, yielding 361 genes expressed in both datasets. The expression profiles in week 4 H9* and H9^NOTCH2NLΔ^ and H9 Week 3, Week 4 and Week 5 were sorted from high to low, and ranked 1 to 361. Then, pairwise comparisons were made between each sample to calculate Spearman’s rank correlation between all samples, and plotted using multi-experiment viewer. 212 genes showed shift towards better correlation with Week 5 data in H9^NOTCH2NLΔ^ compared to H9*. These 212 genes were subjected to GO analysis using Panther V13.0 (Mi et al., 2017). A selection of genes from the significantly associated term neuron differentiation was plotted in a heatmap. Z-scores were calculated for the different samples of Week 4 H9* and H9^NOTCH2NLΔ^, and H9 Week 3, Week 4 and Week 5.

### Co-immunoprecipitation and immunoblot

HEK293T cells were cultured according to standard protocol in DMEM, 4.5 g/l glucose + GlutaMax, 10% HIFBS and 1x P/S. For co-IP experiments, HEK293T cells were transfected in T25 flasks at 50% confluency. pCIG-NOTCH2-Myc and pCAG-NOTCH2NL-HASh + pCAG-NOTCH2NL-HAL,T 197I were mixed in equimolar ratios and transfected using Lipofectamine 2000 (Thermofisher). For control conditions, pCIG-EV and pCAG-EV were used in equimolar ratios. 6 hours after transfection, medium was replaced, and another 24 hours later medium was replaced. Cells were harvested 48 hours after transfection. Cells were washed 3 times with cold 1x PBS, then incubated in 40 minutes in IP buffer (50mM Tris-HCl, 150mM NaCl, 5mM MgCl, 0.5mM EDTA, 0.2% NP-40, 5% glycerol, supplemented with cOmplete, EDTA-free protease inhibitor cocktail (Sigma)) Cells were lysed by passing cell suspension through 273/4 gauge needle 10 times. Lysate was centrifuged 10 minutes at 4°C, supernatant was transferred to a fresh 1.5ml tube. 2 pg of one specific antibody was added (anti-HA Abcam ab9110, anti-Myc Abcam ab9E10, anti-His Abcam ab9108, anti-NOTCH2 SCBT sc25-255) and incubated overnight at 4°C in a rotating wheel. DynaBeads were blocked using 3 washed of 1x PBS + 0.5% BSA and added to IP samples, incubating 3 hours at 4°C rotating. Using a magnetic separator, samples were washed 2 times in cold IP buffer. Then samples were eluted in Tris-EDTA buffer and transferred to new 1.5ml tubes. 2x Laemmli buffer + DTT was added 1:1 prior to SDS PAGE. Samples were loaded on 4-20% Tris glycine gels (Bio-Rad), followed by blotting on nitrocellulose membranes following manufacturer’s recommended protocol. Membranes were blocked in 5% skim-milk powder in 1x PBS + 0.05% Tween or 1x TBS + 0.1% Tween. Primary antibodies were incubated 3 hours at room temperature in 1x PBS (anti-NOTCH2 sc25-255) or 1x TBS-T (other antibodies), followed by 3 washes in 1x PBS-T (anti-NOTCH2 sc25-255) or 1x TBS-T (other antibodies). Secondary antibodies (anti-Rabbit-HRP, anti-Mouse-HRP, Thermofisher) were incubated 60 minutes at room temperature, followed by 3 more washes in 1x PBS-T or 1x TBS-T. Membranes were incubated with supersignal westdura ECL substrate (Thermofisher) and imaged using Bio-Rad Chemidoc imager. For experiments with pCAG-NOTCH2NL-His, pCIG-NOTCH2-Myc, pCIG-PDGFRB-Myc and pCIG-EGFR-Myc, the same protocol was used with equimolar mixes of plasmid DNA. For immunoprecipitation of NOTCH2NLSh,T197I from mouse 46c ESCs, the same protocol was used and protein was isolated from medium using the NOTCH2 sc25-255 antibody.

### NOTCH reporter cell line culture

U2OS cells were cultured in DMEM, 4.5 g/l glucose + GlutaMax, 10% HIFBS and 1x P/S. U2OS-JAG2 cells (gift of Arjan Groot and Marc Vooijs, MAASTRO lab, Maastricht University) were supplemented with 2 μg/ml puromycin. OP9 cells and OP9-DLL1 (gift of Bianca Blom, Academic Medical Center Amsterdam) were cultured in MEMα without nucleosides (Thermofisher), 2mM L-glutamine, 20% HIFBS, 100 μM 2-mercaptoethanol and 1× P/S. For routine culturing, cells were passaged every 3-4 days using 0.25% Trypsin (Thermofisher) + 0.5 mM EDTA (Sigma) in PBS at densities of 1/8 to 1/10 (U2OS), or 1/4 to 1/6 (OP9).

### NOTCH reporter co-culture assays

U2OS cells were seeded at a density of 425,000 cells per well for transfection (6-wells plate). In parallel, U2OS control or U2OS-JAG2 cells were seeded at a density of 110,000 cells per well for co-culture (12-wells plate). After 24 hours, U2OS cells in 6-wells plates were transfected the following amounts of plasmid DNA per well. For control conditions: ng pGL3-UAS, 33.3 ng CMV-Renilla, 16.7 ng pCAG-GFP, 200 ng pcDNA5.1-NOTCH2-GAL4, 167 ng pCAG-EV, and 273 ng pBluescript. For conditions including NOTCH2NL:500 ng pGL3-UAS, 33.3 ng CMV-Renilla, 16.7 ng pCAG-GFP, 200 ng pcDNA5.1-NOTCH2-GAL4, 200 ng pCAG-NOTCH2NL, and 240 ng pBluescript. Plasmid DNA mix was transfected using polyethylenimine (PEI, linear, MW 25000, Polysciences). All amounts were scaled accordingly for multiple transfections. For larger experiments, cells were seeded and transfected in T25 flasks or on 100 mm plates and amounts used were scaled accordingly to surface area. 6 hours after transfection, 6-wells plates were treated with 0.5 ml of 0.25% Trypsin and 0.5 mM EDTA in PBS per well for 2 minutes at 37 degrees. Cells were resuspended in a total volume of 7 ml after addition of culture medium. Medium of 12-wells plates was removed, and 1 ml of transfected cell suspension was added to each well for co-culture. 10 μM DBZ was added to selected control wells. After 24 hours, medium was removed and cells washed once with PBS. Cells were incubated in 150 μl of 1x passive lysis buffer (PLB, Promega) on an orbital shaker for 15 minutes. Lysates were stored at −80°C until analysis. In OP9 and OP9-DLL1 co-cultures, 80,000 cells were seeded per well of a 12-wells plate. For generating conditioned medium, U2OS cells were seeded on 100 mm plates, and were PEI transfected with 2000 ng of pCIG-EV, or 2400 ng of NOTCH2NLA or NOTCH2NLB. Another 10000 ng or 9600 ng of pBluescript was used as carrier DNA. 6 hours after transfection, medium was replaced. 32 hours after transfection, medium was collected and 0.2 μm filtered and used the same day. The experiments using conditioned medium were done as previously described, but were seeded on 0.25% gelatin, 0.1% BSA coated plates instead. For the reporter U2OS cell transfection, only pCAG-EV, and NOTCH2NL plasmids were not added to the plasmid DNA mix, and replaced by pBluescript. Instead, transfected cells are resuspended and seeded in conditioned medium harvested from other cells. For DLL4 assays, 24-wells plates were coated overnight at 4°C with 150 μl of 5 μg / ml rDLL4 (R&D Systems), 0.25% gelatin, 0.1% BSA in PBS. Control plates were coated with 0.25% gelatin, 0.1% BSA in PBS only. U2OS cells were transfected and seeded according to co-culture protocol as previously described, except 0.5 ml of cell suspension was used for each well of the coated 24-wells plates. NOTCH-GAL4 and reporter constructs were kindly gifted by Arjan Groot and Marc Vooijs (MAASTRO lab, Maastricht University).

### RT-PCR characterization of primate *NOTCH2NL* fusion genes

For amplification and detection of potential fusion transcripts, Qiagen one-step RT-PCR kit was used according to manufacturer’s protocol. 25 ng of total RNA isolated from gorilla iPSCs, chimpanzee iPSCs, or human H9 ESCs was used per reaction. Primers used in these reactions were:

Human_N2NL_Fw1_exon1: CGCTGGGCTTCGGAGCGTAG

Human_N2NL_Fw2_exon2: AGTGTCGAGATGGCTATGAA

Human_N2NL_Fw3_exon3: ATCGAGACCCCTGTGAGAAGA

Human_N2NL_Rv2_exon5: CCAGTGTCTAATTCTCATCG

PDE4DIP_Fw1_exon27: AAGGCCCAGCTGCAGAATGC

PDE4DIP_Fw2_exon24: ACACCATGCTGAGCCTTTGC

Chimp_N2NL_Rv1_exon3: GCAAGGTCGAGACACAGAGC

MAGI3_Fw1_exon1: GGGTTCGGGATGTCGAAGAC

MAGI3_Fw2_exon10: GCAACTGTGTCCTCGGTCAC

MAG 13_Fw3_exo n14: GGGAG CAGCTGAGAAAGATG

TXNIP_Fw1_exon1: CAGTTCATCATGGTGATG

TXNIP_Fw2_exon1: GGGTACTTCAATACCTTGCATG

BRD9_Fw1_exon12: GCAGGAGTTTGTGAAGGATGC

BRD9_Fw2_exon10: ACGCTGGGCTTCAAAGACG

### 10× Library Enrichment

To enrich whole-genome sequencing libraries to allow for cost-effective deep sequencing of the *NOTCH2NL* loci, a MYcroarray MyBaits custom oligonucleotide library was developed. 100 bp probes were designed spaced 50 bp apart in chr1:145,750,000-149,950,000, ignoring repeat masked bases, for a total of 20,684 probes. A further 8,728 probes were created in the three *NOTCH2NL* loci by tiling with 50 bp overlaps, ignoring repeat masking but dropping any probes with very low complexity. 17,866 probes were added at every SUN position tiling at 5 bp intervals from −75 bp to +75 bp around the SUN. To try and capture population diversity and ensure even enrichment, at every SNP in the NA12878 Genome In a Bottle variant call set the reference base was replaced and probes tiled in the same fashion as the SUNs. Finally, to reach the required 60,060 probes a random 347 probes were dropped.

### Library Preparation and Enrichment of 10x Chromium Libraries

High molecular weight DNA was processed into Illumina sequencing libraries using the Chromium Genome Reagent Kit V2 chemistry and enriched using the custom MyBaits oligonucleotide probes described above (**Figure S1**). Briefly, high molecular weight (HMW) gDNA was isolated from cultured cells using a MagAttract kit (Qiagen) followed by quantification with Qubit. HMW DNA was partitioned inside of an emulsion droplet along with DNA barcode containing gel beads and an amplification reaction mixture. After barcoding the molecules within the emulsion, Illumina sequencing adaptors were added by ligation. In preparation for hybridization with MyBaits probes Illumina adaptor sequences are blocked with complementary oligonucleotides. Biotinylated probes were hybridized overnight at 65C and isolated using streptavidin coated MyOne C1 beads (Invitrogen). The final enriched libraries were amplified using an Illumina Library Amplification Kit (Kapa).

### Sequencing of Enriched 10x Libraries

The MYcroarray probes were used to enriched 10x Genomics sequencing libraries for three well studied individuals (NA19240, NA12877 and CHM1), the H9 ESC line, the six Simons VIP samples in Figure 7, and the H9 CRISPR mutant in Figure 5. NA12877 was chosen instead of NA12878 because of the existence of high depth 10x whole-genome data for that individual (see below). We find that around 50% of our reads map to regions of enrichment, leading to >1000x coverage of the *NOTCH2NL* loci. The NA19240, NA12877 and H9 libraries were sequenced to 65 million reads, 71 million reads, and 107 million reads respectively. The Simons VIP samples SV721, SV877, SV7720, SV780, SV735 and SV788 were sequenced to a depth of 57 million, 30 million, 44 million, 37 million, 86 million and 37 million reads respectively.

### NOTCH2NL Simons Samples Coverage Analysis

To assess copy number change in the Simons VIP 1q21.1 collection, the H9, NA12877 and NA19240 normal libraries described above were mapped to GRCh38 using Longranger 2.1.3. bamCoverage was used to extract all reads that mapped to the region chr1:142785299-150598866, normalizing depth to 1x coverage across the region to account for library depth. Wiggletools mean (Zerbino et al., 2014) was used to average the depth across these samples. Wiggletools was then used to perform a ratio of this average with the coverages of every Simons 1q21.1 collection sample, which simultaneously normalizes out bias from the array enrichment as well as GC content. These coverages were then re-scaled by the average coverage in the region chr1:149,578,286-149,829,369, which is downstream of *NOTCH2NLC* and not observed to have copy number change. This rescaling adjusts for a systematic shift downward caused by the combination of the previous normalizations seen in deletion samples, and a similar shift upward in duplication samples. Finally, sliding midpoint smoothing was applied to each coverage track, taking into account missing data by ignoring it and expanding the window size symmetrically around a midpoint to always include 100,000 datapoints, stepping the midpoint 10 kb each time.

### Hominid Copy Number Analysis

Sequencing data for NA12878 (ERR194147), Vindjia Neanderthal (PRJEB21157), Altai Neanderthal (PRJEB1265), Denisovan (ERP001519), Chimpanzee (SRP012268), Gorilla (PRJEB2590) and Orangutan (SRR748005) were obtained either from SRA or from collaborators. These data were mapped to GRCh37 to obtain reads mapping to the *NOTCH2* (chr1:120,392,936-120,744,537) and *NOTCH2NL* (chr1:145,117,638-145,295,356) loci in that assembly, and then those reads were remapped to a reference containing just the GRCh38 version of *NOTCH2*. Coverage was extracted with bamCoverage, normalizing to 1x coverage across the custom *NOTCH2* reference. The resulting coverage tracks were then scaled to the average of the unique region of *NOTCH2* then underwent the same sliding midpoint normalization described above, with 5,000 datapoints per window and 2.5 kb step size.

### Gordian Assembler

The extremely low number of long fragments per partition in the 10X Chromium process ensures that nearly all partitions containing sequence from a *NOTCH2NL* repeat will contain sequence from precisely one repeat copy. In order to recover the precise *NOTCH2NL* repeat sequences, a process was developed to assemble paratypes using barcoded reads. A 208 kb multiple sequence alignment of *NOTCH2NL* paralogs was constructed and a consensus sequence generated. For each sample being assembled, the 10x Longranger pipeline was used to map enriched or unenriched reads to GRCh38. All reads that mapped to any of the five *NOTCH2* or *NOTCH2NL* loci in that alignment were extracted, as well as any reads associated with the same input molecules via the associated barcodes. These reads were then remapped to the consensus sequence. FreeBayes^73^ [REF: http://arxiv.org/abs/1207.3907] was used to call variants on these alignments with ploidy set to 10 based on the putative number of NOTCH2NL repeats. Each barcode is then genotyped to find the set of alleles supported at each informative SNP site. Alleles for the majority of SNP sites are undetermined in each barcode due to the sparsity of the linked reads. The result is an MxB sparse matrix where M is the number of variants and B is the number of barcodes identified as having NOTCH2-like sequence. A statistical model is then used to phase this matrix into K paratypes. For each cluster of barcodes representing a single paratype, all reads with the associated barcodes are pooled for short-read assembly using the DeBruijn graph assembler idba_ud (Peng et al., 2012). This software is available on github at http://github.com/abishara/gordian_assembler.

### Establishment of Paratypes in Population

The paratype assembly process described above was applied to the MYcroarray enriched 10x sequencing of NA19240, H9, NA12877, and the six Simons VIP samples. The H9 paratypes were validated with full-length cDNA sequencing (below). The NA12878/NA12891/NA12892 trio (Utah) as well as the NA24385/NA24143/NA24149 (Ashkenazi) trio were assembled using linked read data produced by 10x Genomics for the Genome In A Bottle Consortium. Inheritance was established for the Ashkenazi trio, as well as for the three NA12878 paratypes that assembled. Inherited paratypes are not double counted in **Table S1**. A scaffolding process using alignments of contigs to GRCh38 was performed to construct full-length *NOTCH2NL* loci for each of these assemblies. The *NOTCH2NL* transcripts were annotated and assessed for their protein level features.

### Enrichment and Sequencing of Full-Length cDNA

Full-length cDNA was constructed from both week 5 cortical organoids as well as undifferentiated H9 hESC total RNA similar to previously described protocols (Byrne et al., 2017)] and were enriched using the same MyBaits oligonucleotide set as the 10x Chromium libraries. These cDNA libraries were prepared and sequenced on the Oxford Nanopore MinION. 47,391 reads were obtained for the undifferentiated cells and 118,545 reads for the differentiated cells. The reads were base called with Metrichor. After pooling these datasets, the reads were aligned to GRCh38 to identify putative *NOTCH2NL* reads. 2,566 reads were identified in the week 5 dataset that mapped to *NOTCH2NL,* and 363 in the undifferentiated. Both datasets were filtered for full-length transcripts by requiring at least 70% coverage to the first 1.1 kb of the consensus sequence. This filtering process removed *NOTCH2* like transcripts, leaving a final set of 1,484 transcripts pooled across both timepoints to be analyzed.

### Validating H9 Haplotypes Using Full-Length cDNA

The 1,484 *NOTCH2NL* transcript sequences identified above were aligned to a consensus sequence of H9 ESC transcript paratypes using MarginAlign (Jain et al., 2015). The reads were then reduced into feature vectors containing variant sites along the first 1.1 kb of the consensus to eliminate noise related to alternative transcription stop sites. The feature vectors were aligned using a Hidden Markov Model with one path for each of the paratype assemblies. Since the transcripts are already aligned to a consensus, there is no need for reverse transitions in the model, and since variation or recombination between paralogs is already accounted for in the assemblies, no transitions between paths are allowed. This vastly simplifies the Forward algorithm, and the maximum probability path (usually determined with the Viterbi Algorithm) is trivial to calculate under these conditions. All mismatches were assumed to be errors and were given an emission probability of 0.1 to approximate the error rate of the nanopore. The paratype assembly was validated by showing that there were no recurrent feature vectors that did not align well to any path through this model.

### CRISPR Mutation of *NOTCH2NL* in the H9 ES Line

H9 hESC (WA09, WiCell Research Institute) at passage 42 were plated on a 6-well dish at 40-50% confluency. After 24 hours, cells were treated with 10μM ROCK inhibitor (Y27632; ATCC, ACS-3030) for 1 hour. 2.5μg of each guide plasmid (E2.1& E5.2, Fig. S5 cloned into pX458, Addgene) was then introduced for 4 hours using Xfect DNA transfection reagent (Clontech, 631317). Each guide set was introduced to all 6 wells of a 6-well plate. 48 hours after transfection, cells were dissociated from wells using Accutase cell dissociation enzyme (eBioscience, 00-4555-56), then rinsed twice in PBS supplemented with 0.2mM EDTA, 2% KnockOut Serum Replacement (Thermo Fisher, 10828028), 1% Penicillin-Streptomycin (LifeTech, 15140122), and 2μM thiazovivin (Tocris, 1226056-71-8), and resuspended in a final volume of 1 mL of sorting buffer. The cells were then filtered in a 70μm filter and sorted on a FACS Aria II (BD Biosciences) with a 100pm nozzle at 20psi to select for cells expressing the Cas9-2A-GFP encoded on pX458. Gating was optimized for specificity. Single cells positive for GFP were plated on a 10 cm plate containing 1.5x106 mouse embryonic fibroblasts (MEFs) and cultured in E8 Flex with 2μM thiazovivin (Tocris, 1226056-71-8) for added for the first 24 hours. After growing 5-7 days, individual colonies were manually isolated into 1 well of a 6-well dish on 250,000 MEFs in E8 Flex. 3-5 days later, 3-7 good colonies at passage 42+3 were frozen in BAMBANKER (Fisher Scientific, NC9582225). Remaining cells on MEFs were used for PCR deletion assay. For all subsequent analysis, cells were adapted to culturing on vitronectin (Thermo Fisher A14700) in Gibco’s Essential 8 Flex medium (Thermo Fisher, A2858501).

PCR assay for CRIPSR deletion: For each clone, gDNA was isolated from one 70% confluent well of a 6-well dish using Zymo Quick-gDNA Miniprep kit (Zymo, D3006) according to the manufacturer’s protocol. PCR was performed using approximately 70ng gDNA with Herculase II fusion DNA polymerase (Agilent, 6006745) using primers N2NL E2del_F (5’ CACAGCCTTCCTCAAACAAA 3’) and N2NL E5del_R (5’ GTGCCACGCATAGTCTCTCA 3’). PCR products of the expected size were cloned and sequenced to determine that at least one of NOTCH2NL harbored the expected deletion. Positive clones were subject Chromium library preparation, target enrichment, Illumina sequencing and NOTCH2NL gene assembly as described above.

### Estimate of *NOTCH2* and *NOTCH2NL* Expression in human fetal brain scRNA-Seq data

To asses *NOTCH2NL* expression in the developing brain, we re-analyzed single cell RNA sequencing data from (Nowakowski, et al, *Science,* In Press). Initial analysis of this data showed low expression of *NOTCH2* and *NOTCH2NL* presumably due to removal of multi-mapping reads. To address this, we constructed a custom Kallisto reference based off GENCODE V19 (hg19) where we removed the transcripts ENST00000468030.1, ENST00000344859.3 and ENST00000369340.3. The reads for 3,466 single cells were then quantified against this Kallisto index, and the *NOTCH2 and NOTCH2NL* rows of the resulting gene-cell matrix compared to previously generated tSNE clusters.

### Copy Number Estimates of *NOTCH2NL* in Human Population

The copy number of *NOTCH2NLR* and *NOTCH2NLC* in the human population were established by extracting reads that map to *NOTCH2* (chr1:119,989,248-120,190,000), *NOTCH2NLR* (chrl: 120,705,669-120,801,220), *NOTCH2NLA* (chr1:146,149,145-146,328,264), *NOTCH2NLB* (chrl: 148,600,079-148,801,427) and *NOTCH2NLC* (chr1:149,328,818-149,471,561) from 266 individuals in the Simons Diversity Panel. These reads were then remapped to the 101,143 bp consensus sequence of a multiple sequence alignment of alignable portions of *NOTCH2* and all *NOTCH2NL* paralogs. This multiple sequence alignment was used to define our SUN markers, and the ratio of reads containing a SUN to a non-SUN were measured and the median value taken for *NOTCH2NLC* and *NOTCH2NLR.* Establishing copy number with SUNs proved difficult for *NOTCH2NLA* and *NOTCH2NLB* due to the high rate of segregating ectopic gene conversion alleles in the population. Each of the 266 samples was studied by hand. Using comparison to the 10 normal genomes assembled, it appeared that *NOTCH2NLA* and *NOTCH2NLB* are not copy number variable in the phenotypically normal population.

### Paratype Estimation of *NOTCH2NL* in Human Population

Assigning paratypes without assemblies is not possible. To try and evaluate the gene conversion landscape in the population, we took the ratio of SUN read depths in all 266 Simons individuals as well as the six Simons VIP samples and our 10 assembled genomes and plotted them split up by paralog (**Table S4**). These were evaluated for *NOTCH2NLC* and *NOTCH2NLR* copy number (**Figure S1G**). Three samples were identified in Simons with apparent gene conversion in *NOTCH2NLC*, which we did not observe in any of our assembled samples. Manual analysis of these SUN diagrams led to the annotation of six distinct classes of *NOTCH2NLA-NOTCH2NLB* gene conversion with varying population frequencies. In some cases, the data were of lower quality and harder to interpret. The most common gene conversion allele is an overwrite of around 20kb of *NOTCH2NLB* by *NOTCH2NLA* in intronic sequence between exons 2 and 3, present in 42.5% of Simons normals haplotypes. When interpreting these SUN plots, it is helpful to remember that the denominator of the ratio is the total copy number, and as such as individuals stray from N=10 the expected values change. Gene conversion can be observed as regions where one paralog has ratios on the Y axis go up while the other goes down. Exons 1-5 are located at 19,212-19,590 bp, 59,719-59,800 bp, 84,150-84,409 bp, 92,421-92,756 bp and 93,009-97,333 bp respectively in the consensus sequence.

### Copy Number Estimates of Microarray 1q21.1 Deletion/Duplication Syndrome Patients

Comparative genomic hybridization (CGH) microarray probes from Agilent array designs #014693, #014950, #24616, and #16267 were mapped to the human hydatidiform mole genome assembly using the pblat aligner (http://code.google.com/pZpblat/). IDs for microcephaly and macrocephaly patients visualized in **Figure S7** are as follows: control-1: MCL08273; control-2: GSM1082800; 1: LG_252808110380_S01; 2: WSX002375; 3: SGM250214; 4: MCL08277; 5:MUG249341; 6: MCL04601; 7: MCL00270; 8: MCL02135; 9: NGS260131; 10: MCL01089; 11: MCL01415. The relative affinities of each probe for each of its mapping locations were calculated using the DECIPHER R package; the predicted hybridization efficiency of a probe and a mapping location was divided by the predicted hybridization efficiency of the probe and its reverse complement, yielding the affinity for the mapping location. An integer linear programming (ILP) model was created for each sample, with integral variables representing the copy number of each probe mapping location in the CGH sample genome and the CGH reference genome, and additional variables representing the total affinity-weighted copy number of all mapping locations for each probe. Approximate equality constraints were added to represent the measured sample/normal hybridization ratios (normalized per-probe to an average control value of 1.0 for arrays with control samples (GEO accession GSE44300) available), the chromosomal structure of the genome, and the prior belief that the copy number at most locations would be 2. Each approximate equality constraint was constructed by creating two variables, which were restricted to be positive and greater than the difference between the constrained quantities in each direction; the weighted sum of these two variables was then added into the model's objective function. In an alternate method, the locations of homologous sequences that could provide a plausible mechanism for duplication or deletion of a region were also added (data not shown). Finally, the resulting ILP model was minimized using CBC^79^ (http://projects.coin-or.org/) to produce the sample's integral copy number calls. The copy number inference software used is available from http://bitbucket.org/adamnovak/copynumber.

### NOTCH2NL Expression in Week 5 Neurospheres

Two replicates of bulk RNA-seq of week 5 cortical organoids derived from H9 ES as well as undifferentiated cells from the H9 differentiation time course described above were quantified against a custom Kallisto reference based on GENCODE V27. Using bedtools, all transcripts which overlapped our curated annotations of *NOTCH2NL* paralogs and *NOTCH2* were removed. After converting this annotation set to FASTA, a subset of our paratype assemblies of H9 *NOTCH2NL* paralogs were added in. Only one representative of both *NOTCH2NLR* and *NOTCH2NLC* was used due to their high similarity on the transcript level. The TPM values of the replicates were averaged.

